# Intracellular localisation of *Mycobacterium tuberculosis* affects antibiotic efficacy

**DOI:** 10.1101/2020.11.25.398636

**Authors:** Pierre Santucci, Daniel J. Greenwood, Antony Fearns, Kai Chen, Haibo Jiang, Maximiliano G. Gutierrez

## Abstract

To be effective, chemotherapy against tuberculosis (TB) must kill the intracellular population of *Mycobacterium tuberculosis* (Mtb). However, how host cell environments affect antibiotic accumulation and efficacy remains elusive. Pyrazinamide (PZA) is a key antibiotic against TB, yet its behaviour is not fully understood. Here, by using correlative light, electron, and ion microscopy to image PZA at the subcellular level, we investigated how human macrophage environments affect PZA activity. We discovered that PZA accumulates heterogeneously between individual bacteria in multiple host cell environments. Crucially, Mtb phagosomal localisation and acidification increase PZA accumulation and efficacy. By imaging two antibiotics commonly used in combined TB therapy, we showed that bedaquiline (BDQ) significantly enhances PZA accumulation by a host cell mediated mechanism. Thus, intracellular localisation and specific microenvironments affect PZA accumulation and efficacy; explaining the potent *in vivo* efficacy compared to its modest *in vitro* activity and the critical contribution to TB combination chemotherapy.

## Introduction

*Mycobacterium tuberculosis* (Mtb), the etiologic agent of tuberculosis (TB), is the most prevalent cause of mortality due to an infectious agent worldwide ^1^. Drug-sensitive TB requires treatment with a minimum of four antibiotics over a course of at least 6 months ^1^. The standard first-line multidrug therapy includes the drugs isoniazid (INH), rifampicin (RIF), ethambutol (ETB) and pyrazinamide (PZA). The duration and toxicity of the current anti-TB regimens affect compliance, leading to treatment failure, relapse and emergence of resistance. The increasing number of multidrug-resistant strains constitutes a global health issue, and new therapeutic strategies are needed to reduce the treatment duration of drug-sensitive TB. However, this is challenging in the context of TB chemotherapy, since the environments within tuberculous granulomas are heterogeneous and dynamic^2,3^.

Mtb is a pathogen able to infect, adapt, survive and replicate in several cell types within its human host. During pulmonary infection, Mtb faces multiple environments, where alveolar macrophages constitute a key replication niche in early infection and thus play a central role in the tubercle bacilli lifecycle ^4^. The intracellular lifestyle of Mtb represents a crucial stage in TB pathogenesis and it is now accepted that drug discovery programmes should include *in cellulo* studies using infected host cells^5,6,7,8^; which may be more accurate than classical *in vitro*-based screens using only isolated bacteria. Within macrophages, Mtb encounters diverse subcellular environments including both membrane-bound compartments, such as phagosomes, phagolysosomes and autophagic compartments, and the cytosol ^9,10,11^. These environments exhibit distinct biophysical and biochemical properties such as nutrient availability, pH, and hydrolytic activities that can affect Mtb replication ^12,13^. In this context, macrophage activation enhances anti-mycobacterial activities and changes Mtb sensitivity towards first-line drugs ^14^. Despite its physiological relevance, if bacterial compartmentalisation alters drug accumulation and efficacy remains poorly characterised. Moreover, how specific subcellular microenvironments impact antibiotic mode of action is elusive.

The contribution of intracellular environments to antibiotic efficacy is particularly important for antitubercular compounds such as the front-line drug PZA; that is mainly effective *in vivo* but has very little potency *in vitro* ^15,16,17,18^. The activity of PZA against Mtb was discovered in the 1950s using TB mouse models before being further investigated in drug-susceptible TB patients ^19^. The remarkable efficacy of PZA played a key role in shortening the anti-TB chemotherapy from 9 to 6 months ^20^. PZA synergises with new drugs such as pretomanid or bedaquiline (BDQ) and therefore it is included in new generations of shorter and less toxic regimens ^21,22,23,24^. Some of these combinations supported by the Global Alliance for TB Drug Development are currently under phase III clinical development, including the trials *STAND* and *SimpliciTB* ^25,26^.

Although PZA is widely used, the contrasting differences between its *in vitro* and *in vivo* efficacy is still not fully understood. Pioneering studies demonstrated that PZA is inactive *in vitro* at neutral pH but displays activity against Mtb at pH 5.5 or below ^16,27,28^. PZA is a prodrug that requires conversion by the mycobacterial nicotinamidase/pyrazinamidase PncA into pyrazinoic acid (POA), which is subsequently exported to the extracellular milieu. If this environment is acidic, POA becomes protonated (HPOA) and crosses the bacterial envelope to finally accumulate within Mtb ^29,30^. This pH-dependent model remains a commonly accepted mechanism of PZA accumulation where HPOA is subsequently able to disrupt membrane potential and raise intrabacterial pH ^28,31^. However, PZA pH-independent activity have also been reported and additional PZA/POA modes of action proposed ^32,33,34^. For example, PZA also inhibits the biosynthesis of Coenzyme-A through inhibition of the aspartate decarboxylase PanD and enhancement of its degradation by the Clp protease system ^35,36,37^. Thus, the molecular mechanisms responsible for PZA efficacy are complex and diverse. Notably, most of the mechanistic studies are performed *in vitro* and do not consider the intracellular lifestyle of Mtb.

Here, we used a recently developed correlative light, electron and ion microscopy (CLEIM) approach to visualize antibiotics at a subcellular resolution ^38^ and investigate PZA distribution in human primary macrophages infected with Mtb. By combining high-content microscopy with genetic and pharmacological perturbations, we analysed how changes in Mtb intracellular localisation and host cell environments affect PZA localisation, accumulation and efficacy. We provide evidence that acidic pH is an important factor underlying PZA efficacy but it is not essential *in cellulo*. Finally, by imaging the subcellular distribution of two different anti-TB antibiotics for the first time, we show that BDQ enhances PZA accumulation in intracellular mycobacteria through a host-dependent mechanism. Our results show that efficacy may be affected by the localisation of bacteria at the time of treatment and provide new directions for the design of future antibiotics and combined therapies.

## Results

### PZA/POA localises primarily within intracellular Mtb

To define the subcellular distribution of PZA, human blood monocyte-derived macrophages (MDM) were infected with Mtb and analysed by correlative electron and ion microscopy (**Figure S1**) ^38^. Previous studies showed that the average PZA maximum serum concentration (*C*_max_) in PZA-treated TB patients is circa 30.8 mg/L ^39^. In order to mimic this physiological concentration and better define PZA distribution *in cellulo*, MDM were infected with Mtb for 24 h and further treated with 30 mg/L of [^15^N_2_, ^13^C_2_]-labelled PZA for an additional 24 h. Because both isotopic labels are present on the pyrazine ring, they are preserved when PZA is converted to POA by the bacterial enzyme PncA. Consequently, ion microscopy cannot differentiate between PZA and POA or other metabolites, which will collectively be referred here as PZA/POA. PZA/POA was detectable by the enrichment of ^15^N relative to ^14^N in treated samples (**Figure S1, Figure 1**) but not by the ^13^C label when compared to control samples, indicating that a higher concentration of ^13^C might be required to be detectable by ion microscopy, due to higher natural abundance of ^13^C (1.1%) compared to the natural abundance of ^15^N (0.37%), and ^13^C background signals from the resin. We observed that ^15^N/^14^N ratio was only above natural abundance within Mtb but not in the host cell cytosol or any other macrophage organelles (**Figure 1A**). Interestingly, some infected macrophages were totally devoid of PZA/POA enrichment (**Figure 1B**). Also, in some PZA/POA positive cells the accumulation was highly heterogeneous, even between neighbouring bacteria (**Figure 1A**). This suggested that each individual host macrophage may have effects on the bacterial PZA/POA accumulation. To test this hypothesis, the distribution of PZA/POA enrichment between bacteria within each single macrophage was compared to the distribution of enrichment across the infected macrophage population (**Figure 1C**). Consistent with the hypothesis, the intra-macrophage standard deviation of accumulation was found to be statistically lower than at the whole-population level (**Figure 1C**). Moreover, analysis of PZA/POA enrichment within Mtb contained in intact cells (**Figure 1**) or necrotic cells revealed that plasma membrane integrity positively impacts drug accumulation (**Figure S2**). Given that loss of membrane integrity in necrotic cells dissipates pH gradients, this finding is consistent with a pH-dependent accumulation of PZA. Thus, PZA/POA accumulation in the context of infection is highly heterogenous between host cells and also between intracellular Mtb.

**Figure 1:**
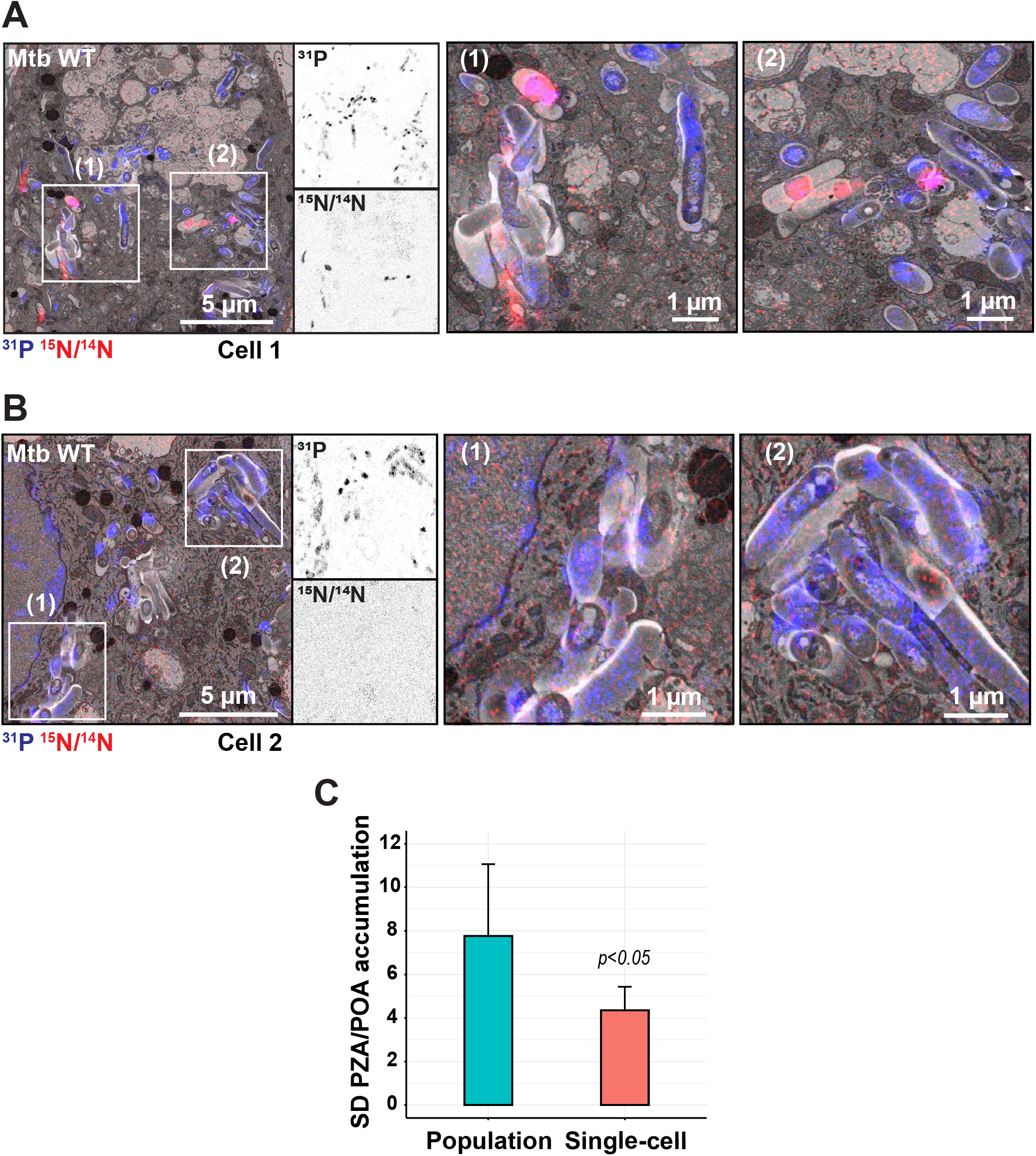
Heterogeneous accumulation of PZA/POA within Mtb-infected MDM. **(A-B)** Mtb-infected MDM treated with 30 mg/L [^15^N_2_, ^13^C_2_]-PZA for 24 hours. EM micrograph is overlaid with ^31^P (blue) and ^15^N/^14^N (red) NanoSIMS images. Magnifications show ^31^P (top panel) and ^15^N/^14^N (bottom panel) individual signals at the single bacterialcell level. Scale bar corresponds to 5 μm. Regions of interest (1) and (2) highlighted by white rectangles, are shown in detail in the right panels respectively. Scale bar corresponds to 1 μm. **(C)** Comparison of standard deviation (SD) in ^15^N/^14^N ratiometric signals between bacteria contained within the same macrophage (single-cell level) and across Mtb in all macrophages (population level). A total of 1659 intracellular bacteria were analysed from 4 biological replicates. Results are expressed as mean of standard deviation +/−confidence interval of 0.95. Statistical significance was assessed by using a t-statistic test from the linear model.

### The PZA/POA accumulation in Mtb is affected by the intracellular localisation

Because low pH has been shown to enhance PZA enrichment and activity under certain culture conditions, we next analysed using ion microscopy whether Mtb subcellular localisation could affect antibiotic distribution and/or accumulation within host cells.

We first used a genetic approach by comparing Mtb WT, which can localise in the cytosol, with the Mtb ΔRD1 mutant that lacks the ESX-1 Type 7 secretion system and resides primarily in membrane-bound compartments ^10,11,40^. We postulated that the restriction of ESX-1 deficient-Mtb to membrane bound compartments would increase the proportion of bacteria in acidic microenvironments, as previously described ^41,42^, and thus might impact PZA/POA accumulation. Co-localisation analysis of Mtb WT and Mtb ΔRD1 infected cells stained with the acidotropic dye LysoTracker confirmed that Mtb ΔRD1 was more often associated with LysoTracker positive acidic environments (**Figure S3**). In addition, to exclude any intrinsic variation in PZA susceptibility between Mtb WT and Mtb ΔRD1, a drug susceptibility assay was performed *in vitro*. Our results were in line with previously published data regarding the weak efficacy of PZA *in vitro* in neutral broth ^27^ and further confirmed that there were no PZA dose-response differences in the relative growth between the two strains (**Figure S4A**). Moreover, no differences in ^15^N/^14^N signal were observed between Mtb WT and Mtb ΔRD1 when incubated for 24 h in the presence of [^15^N_2_, ^13^C_2_]-labelled PZA *in vitro* (**Figure S4B**). A quantitative analysis of PZA/POA accumulation in Mtb-infected macrophages revealed a higher level of antibiotic associated with Mtb ΔRD1 than the Mtb WT strain (average mean intensity per positive bacteria of 93.02 ± 4.98 *vs.* 65.77 ± 3.27, *p*<0.01) (**Figure 2A-C**), consistent with the proposal that membrane-bound compartments promote PZA/POA accumulation. Analysis of the distribution of PZA/POA positive bacteria between Mtb WT and Mtb ΔRD1 by density plot confirms that WT bacteria harbouring PZA/POA positive signature displayed a lower enrichment than ΔRD1 bacteria (**Figure 2E**). In addition to a significant increase in ^15^N/^14^N ratio per bacteria, the percentage of PZA/POA positive bacteria was also higher during infection with the Mtb ΔRD1 strain (**Figure 2D**).

**Figure 2:**
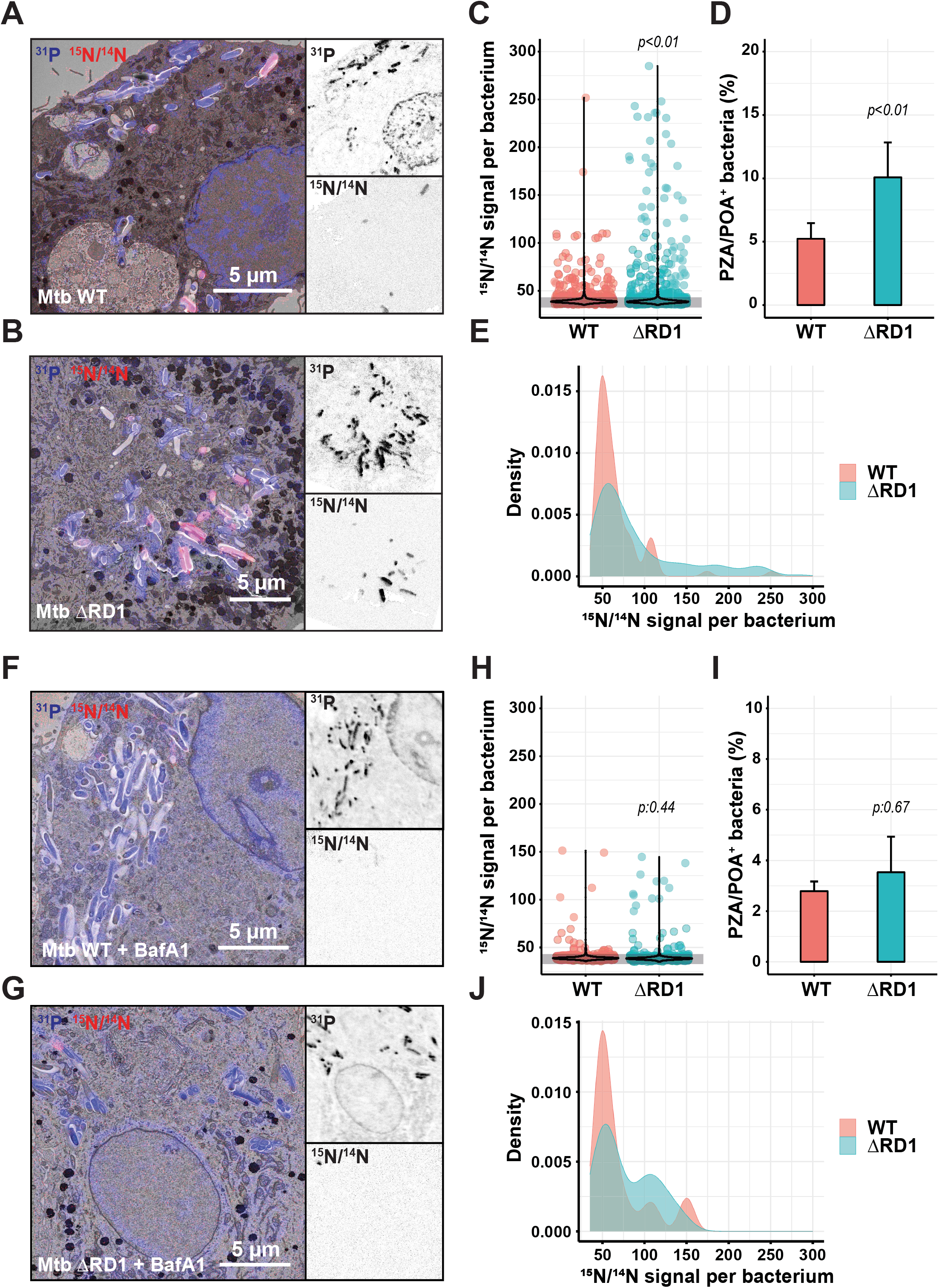
Subcellular localisation and endolysosomal acidification contribute to PZA/POA accumulation in intracellular Mtb. **(A-B)** Representative images of PZA/POA distribution in intracellular Mtb WT and Mtb ΔRD1 strains. MDM were infected with Mtb WT or Mtb ΔRD1 and treated with 30 mg/L [^15^N_2_, ^13^C_2_]-PZA for 24 hours. EM micrographs are overlaid with ^31^P (blue) and ^15^N/^14^N (red) NanoSIMS images. Magnifications show ^31^P (top panel) and ^15^N/^14^N (bottom panel) individual images at the single bacterial-cell level. Scale bars correspond to 5 μm. **(C)** Quantitative analysis of ^15^N/^14^N signal per bacterium shown as violin plot with single dots. Grey line indicates the natural background level of the ^15^N/^14^N enrichment. Results were obtained from 1352-1649 individually segmented bacteria and p-values were calculated from the linear model. **(D)** Quantification of the percentage of PZA/POA positive (PZA/POA^+^) bacteria. Results are expressed as mean ± standard error of the mean from 3-4 biological replicates and p-values were calculated from the linear model. **(E)** Analysis of the ^15^N/^14^N signal profile of Mtb WT and Mtb ΔRD1 PZA/POA positive bacteria. **(F-G)** Representative images of PZA/POA distribution in intracellular Mtb WT and Mtb ΔRD1 strains treated with 30 mg/L [^15^N_2_, ^13^C_2_]-PZA for 24 hours in the presence of 100 nM BafA1. Micrographs and magnifications are displayed as described for **(A-B)**. Scale bars correspond to 5 μm. **(H)** Quantitative analysis of ^15^N/^14^N signal per bacterium shown as violin plot with single dots. Grey line indicates the natural background level of the ^15^N/^14^N enrichment. Results were obtained from 741-828 individually segmented bacteria and p-values were calculated from the linear model. **(I)** Quantification of the percentage of PZA/POA positive (PZA/POA^+^) bacteria after 24 hours of 100 nM BafA1 treatment. Results are expressed as mean ± standard error of the mean from 2-3 biological replicates and p-values were calculated from the linear model. **(J)** Analysis of the ^15^N/^14^N signal profile of Mtb WT and ΔRD1 PZA/POA positive bacteria in the presence of 100 nM BafA1.

Next, we postulated that if the pH-dependent model of PZA accumulation is correct, inhibition of endosomal acidification would result in reduced enrichment of the antibiotic within intracellular bacteria. To test this hypothesis, macrophages were infected with either Mtb WT or Mtb ΔRD1 and further treated with [^15^N_2_, ^13^C_2_]-PZA in the absence or presence of the pharmacological modulator BafA1, a specific inhibitor of the mammalian vacuolar-type H^+^-ATPase ^43^. We first assessed BafA1 activity and confirmed that treatment for 2 h, 24 h or 72 h significantly reduced pH-dependent proteolytic activity of macrophages without affecting cell viability (**Figure S5-S6**). As expected, BafA1 treatment of Mtb-infected cells for 24 hours resulted in a significant diminution of bacteria in acidic compartments (**Figure S3**). The level of PZA/POA in treated macrophages was still heterogenous with few bacteria that strongly accumulated PZA/POA whereas many others were negative for ^15^N/^14^N enrichment (**Figure 2F-G**). Quantitative analysis of intrabacterial ^15^N/^14^N ratios in the presence or absence of BafA1 revealed that inhibition of endolysosomal acidification significantly reduced PZA/POA level in Mtb ΔRD1 (average mean intensity per positive bacteria of 93.02 ± 4.98 *vs* 78.5 ± 5.93, *p*<0.05) but were similar in Mtb WT (average mean intensity per positive bacteria of 65.77 ± 3.27 *vs* 69.19 ± 7.48, *p*:0.36) when compared to their respective untreated control samples (**Figure 2H and 2C**). Moreover, pairwise comparison between Mtb WT and Mtb ΔRD1 upon BafA1 treatment showed no significant differences in mean ^15^N/^14^N signal and percentage of PZA/POA positive bacteria (*p*:0.44 and *p*:0.67, respectively). That this decrease occurs primarily with Mtb ΔRD1 upon endolysosomal acidification inhibition argues that Mtb restricted to membrane-bound compartments are more susceptible to PZA/POA accumulation. Altogether, these results support a pH-dependent model of PZA accumulation within intracellular Mtb, highlighting that the subcellular localisation of the tubercle bacilli and the nature of the microenvironments it encounters within human macrophages impact PZA accumulation.

### The intracellular localisation of Mtb affects PZA efficacy

Next, we investigated if these changes in bacterial intracellular localisation and PZA/POA enrichment had an effect on efficacy. The effect of PZA on intracellular Mtb replication was assessed by using a high content single-cell analysis imaging approach ^38^ (**Figure S7**). After 96 h of infection, Mtb WT replicated faster than the Mtb ΔRD1 mutant in resting human macrophages (**Figure 3A-B**) as previously reported ^10,40^. Replication inhibition profiles observed with Mtb WT at 30-and 100 mg/L agreed with previous results obtained in PZA-treated human macrophages after 96 h of infection showing that PZA is efficient *in cellulo* ^44^. In contrast to our *in vitro* results, Mtb ΔRD1 was more susceptible to PZA than Mtb WT *in cellulo* (**Figure 3A-B**). Indeed, analysis of total bacterial levels at 72 h post-treatment in comparison to the 24 h pre-treatment values demonstrated that PZA was more effective against Mtb ΔRD1 when compared with Mtb WT (**Figure 3B**). A relative growth inhibition analysis showed that at 30 mg/L, PZA is more effective against Mtb ΔRD1 than Mtb WT (**Figure 3E**). Interestingly, at a higher concentration (100 mg/L), the inhibition was more efficient but the difference between the Mtb WT and Mtb ΔRD1 was reduced, suggesting that at higher concentrations PZA might distribute more evenly into more intracellular microenvironments.

**Figure 3:**
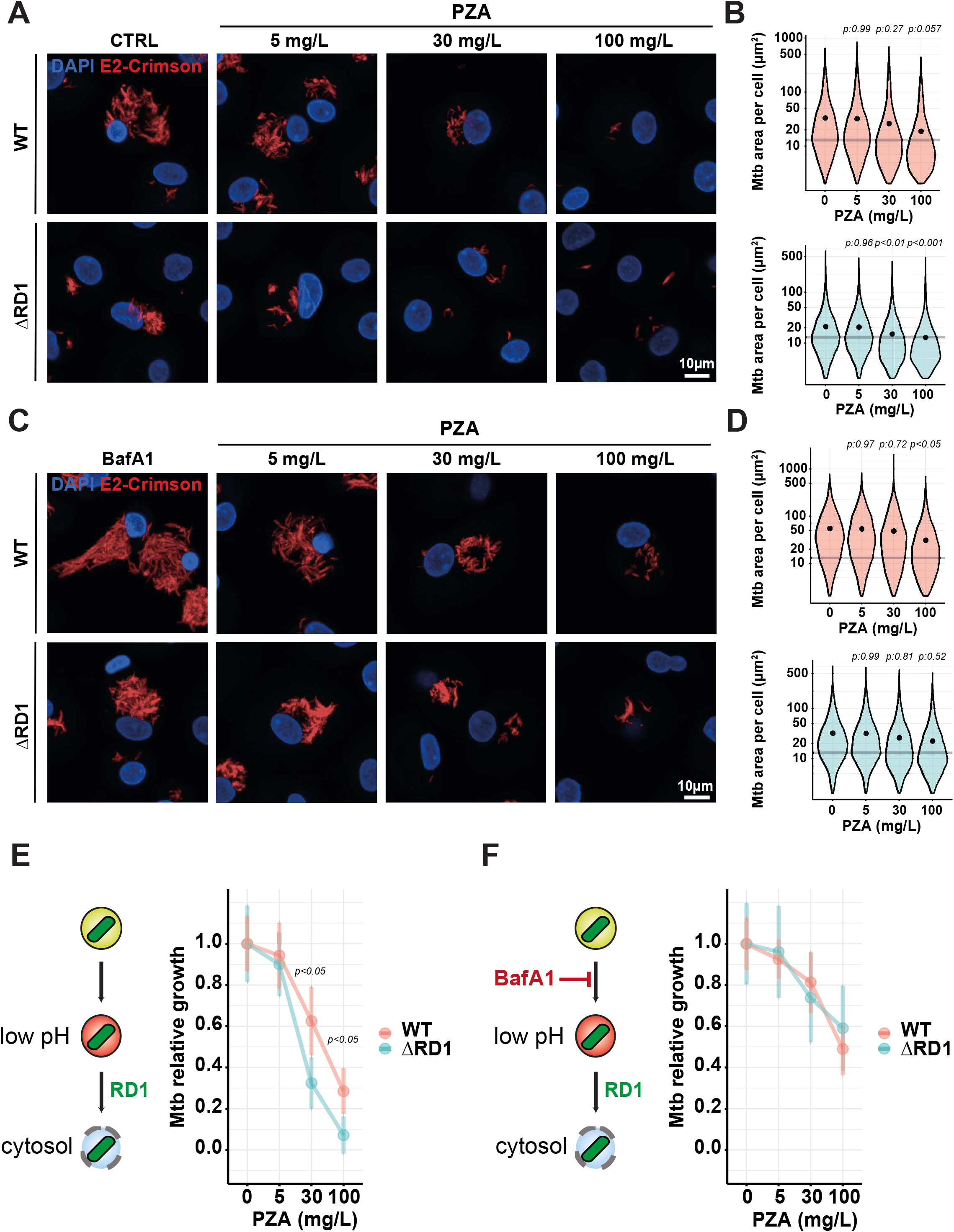
PZA inhibits Mtb replication *in cellulo* and is more effective against Mtb ΔRD1 mutant strain. **(A)** Representative confocal fluorescence images of Mtb WT- and Mtb ΔRD1-infected MDM for 24 hours and further treated for 72 hours with increasing concentration of PZA. Magnifications display nuclear staining (blue) and Mtb-producing E2-Crimson (red). Scale bar corresponds to 10 μm. **(B)** Quantitative analysis of E2-Crimson Mtb WT (top panel) and Mtb ΔRD1 (bottom panel) area per single-cell expressed in μm^2^. Results are displayed in violin plots where grey lines represent the mean Mtb area per cell pretreatment (t24h) and black dots represent the mean Mtb area per cell post-treatment (t_96h_). From 7124 to 8785 and 5419 to 6560 infected MDM were analysed for Mtb WT and Mtb ΔRD1 respectively. Results are from 4 biological replicates and statistical significance was assessed by comparing the means of each conditions using one-way ANOVA followed with Tukey’s multiple comparisons test. **(C)** Representative confocal fluorescence images of Mtb WT- and Mtb ΔRD1-infected MDM for 24 hours and further treated for 72 hours with increasing concentration of PZA in the presence of 100 nM BafA1. Magnifications are displayed as described in **(A)**. **(D)** Quantitative analysis of E2-Crimson Mtb WT (top panel) and Mtb ΔRD1 (bottom panel) area per single-cell expressed in μm^2^, is represented as in **(B)**. From 7821 to 9130 and 6619 to 7723 infected MDM were analysed for Mtb WT and Mtb ΔRD1 respectively. Results are from 4 biological replicates and statistical significance was assessed by comparing the means of each conditions using one-way ANOVA followed with Tukey’s multiple comparisons test. **(E-F)** Mean bacterial area per macrophage in the presence or absence of 100 nM BafA1 and increasing concentration of PZA was normalized and plotted as relative growth. A schematic representation of the BafA1 effect onto endolysosomal pH accompanied the graphs. Normalization was done to mean Mtb area per cell pre-treatment (t_24h_) and the control condition without PZA was used as reference corresponding to 100 % growth. Results are displayed as mean of the 4 biological replicates +/− standard error of the mean and statistical significance was assessed between Mtb WT and Mtb ΔRD1 by comparing the means of each conditions using one-way ANOVA followed with Tukey’s multiple comparisons test.

Then, we tested if modulation of endolysosomal acidification affects PZA efficacy. For this, macrophages were infected with Mtb WT or Mtb ΔRD1 for 24 h and then treated with PZA in presence or absence of the vacuolar-type H^+^-ATPase inhibitor BafA1 (therefore an inhibitor of endolysosomal acidification). Treatment with BafA1 clearly increased both Mtb WT and Mtb ΔRD1 replication in macrophages (**Figure 3C-D and Figure S8**), confirming that acidification/proteolytic activity is mostly restrictive for Mtb at some stages of the intracellular lifestyle ^45^. Similar results were obtained with Concanamycin A (ConA), another vacuolar-type H^+^-ATPase inhibitor ^46^ (**Figure S8**). Neither BafA1 nor ConA had a direct effect on Mtb WT or Mtb ΔRD1 growth *in vitro* (**Figure S8**). BafA1 treatment reduced PZA efficacy when used at 30 mg/L resulting in a significantly higher bacterial burden, with changes in inhibition values from 40% to 20% for Mtb WT and from 70% to 25% for the Mtb ΔRD1 mutant (**Figure 3F**). These differences in efficacy were confirmed by colony-forming units (CFU) analysis (**Figure S9**). Interestingly, the greater effect of BafA1 on Mtb ΔRD1, which are mostly in a membrane bound/acidic environment, supports a role for low pH in PZA efficacy. Notably, although BafA1 greatly reduced PZA efficacy against intracellular bacteria, PZA was still effective against these bacteria and reduced Mtb replication. This argues that an acidic environment may enhance PZA efficacy but it is not strictly required. Importantly, this effect was specific for PZA, since BafA1 treatment did not significantly affect the efficacy of RIF or INH (**Figure S10**). Altogether, these data support the existence of both pH-dependent and independent mechanisms of PZA accumulation and activity against intracellular Mtb, providing evidence that Mtb localisation had a significant impact in PZA efficacy.

### PZA/POA accumulates in Mtb residing in non-acidic compartments

Our analysis suggested that both pH-dependent and pH-independent mechanisms contribute to PZA efficacy *in cellulo*. To define whether distinct intracellular populations of Mtb accumulate PZA/POA in a pH-dependent manner, we used a CLEIM approach. Macrophages were infected with fluorescent Mtb, treated with 30 mg/L of [^15^N_2_, ^13^C_2_]-PZA for 24 h and acidic compartments were labelled using LysoTracker. Within single macrophages, we analysed the PZA/POA enrichment in both LysoTracker positive and LysoTracker negative Mtb (**Figure 4**). We observed that most of the intracellular bacteria analysed within a single macrophage were harbouring extremely low levels of PZA/POA almost similar to the natural background enrichment (^15^N/^14^N signal values ranging from 37 to 44) regardless of the pH microenvironment, as measured by LysoTracker staining. Strikingly, two bacteria contained in a LysoTracker negative compartment accumulated higher levels of PZA/POA, suggesting that PZA/POA enrichment can occur in non-acidic environments (**Figure 4**). Altogether, these data revealed that intrabacterial PZA/POA accumulation within host cells can be pH-independent. These observations might explain the highly heterogeneous enrichment of both Mtb WT and Mtb ΔRD1, but also the partial inhibitory effect observed in the presence of vacuolar-type H^+^-ATPase inhibitor. These results might also explain why PZA is still very effective against Mtb WT despite its localisation in the cytosol.

**Figure 4:**
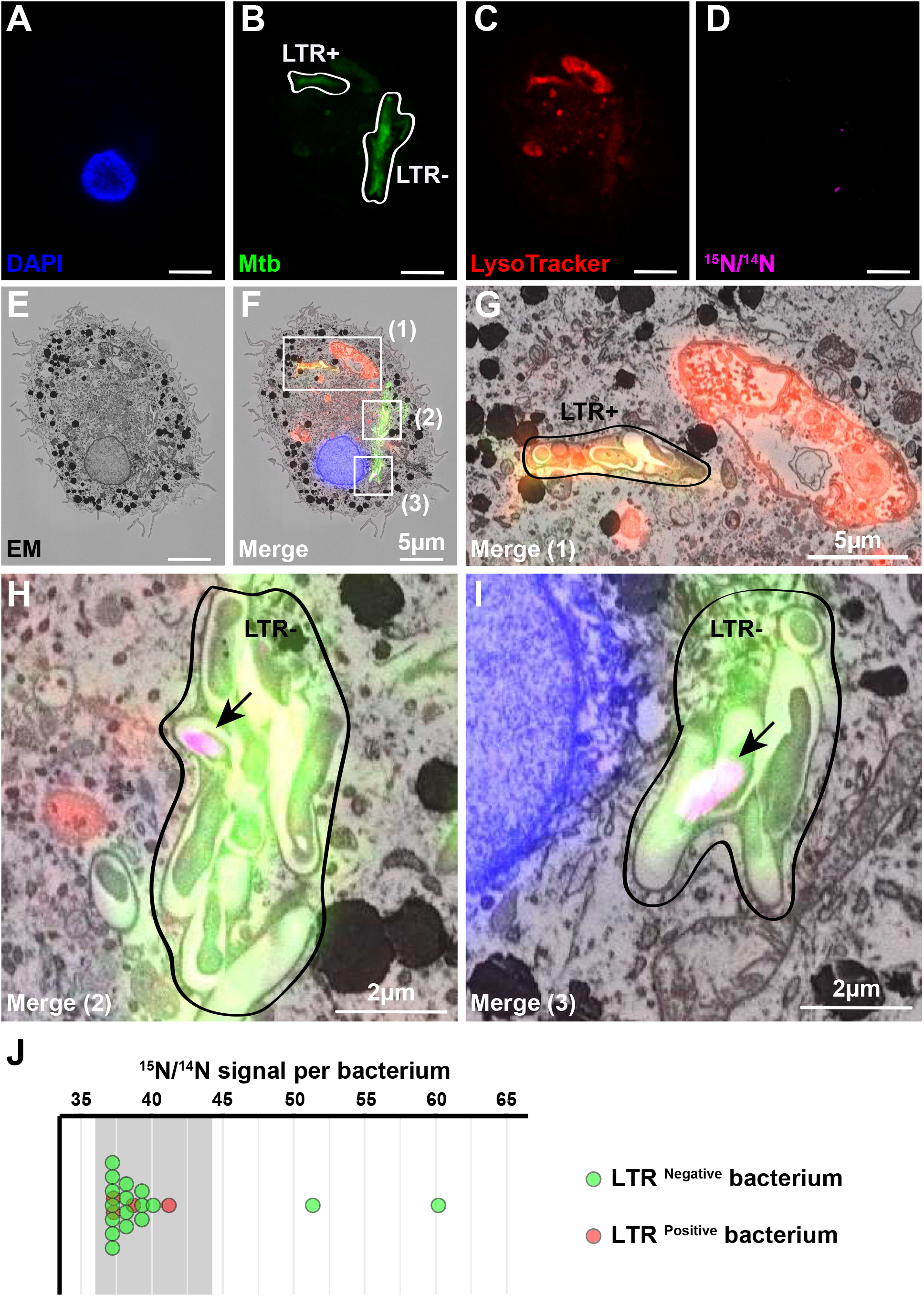
CLEIM reveals that PZA/POA is able to accumulate within Mtb residing in non-acidic compartments. **(A-I)** Mtb-infected MDM were treated with 30 mg/L [^15^N_2_, ^13^C_2_]-PZA for 24 hours. The sample was then stained with 200 nM LysoTracker before chemical fixation and counterstained with DAPI. After confocal fluorescence imaging, sample was embedded and the cell of interest was processed for correlative electron and ion microscopy. Representative fluorescence images corresponding to **(A)** nuclear staining (blue), **(B)** Mtb WT (green), **(C)** LysoTracker (red) were overlaid with **(D)** the ^15^N/^14^N signal (purple) and **(E)** the EM micrograph (black and white). Merging of the five channels is shown in **(F)** where three regions of interest numbered (1), (2) and (3) respectively, are highlighted by white rectangles. Scale bars correspond to 5 μm. **(G)** Magnification of region (1) shows bacteria contain within LTR positive compartment devoid of any detectable PZA/POA. Scale bar corresponds to 5 μm. **(H-I)** Magnifications of regions (2) and (3) highlight PZA/POA accumulation in LTR negative microenvironment. Scale bar corresponds to 2 μm. **(J)** Quantification of ^15^N/^14^N signals associated with each bacterium contained in LysoTracker positive and negative microenvironments. Grey line indicates ^15^N/^14^N natural background level. Bacterium with strong positive ^15^N/^14^N signals correspond to the bacterium highlighted in (2) and (3) respectively.

### BDQ enhances the accumulation of PZA in intracellular Mtb

In macrophages, BDQ enhances the targeting of Mtb to acidic compartments and synergises with PZA ^47^. We hypothesized that BDQ-dependent targeting to acidic compartments will increase PZA enrichment within intracellular Mtb. BDQ did not increase PZA/POA enrichment per bacterium when co-treated at both neutral and acidic pH *in vitro* (**Figure S11**). In contrast, BDQ significantly enhanced PZA/POA accumulation in intracellular Mtb (**Figure 5A-B**). A quantitative analysis of both BDQ and the PZA/POA levels per single-bacterium showed no correlation between the amount of BDQ per bacterium and the increase in PZA/POA (R=0.14, *p*<0.01). This agrees with previous reports suggesting that BDQ triggers host cell-mediated changes that impacts PZA accumulation/efficacy ^23, 47^, rather than through a bacterial-mediated mechanism.

**Figure 5:**
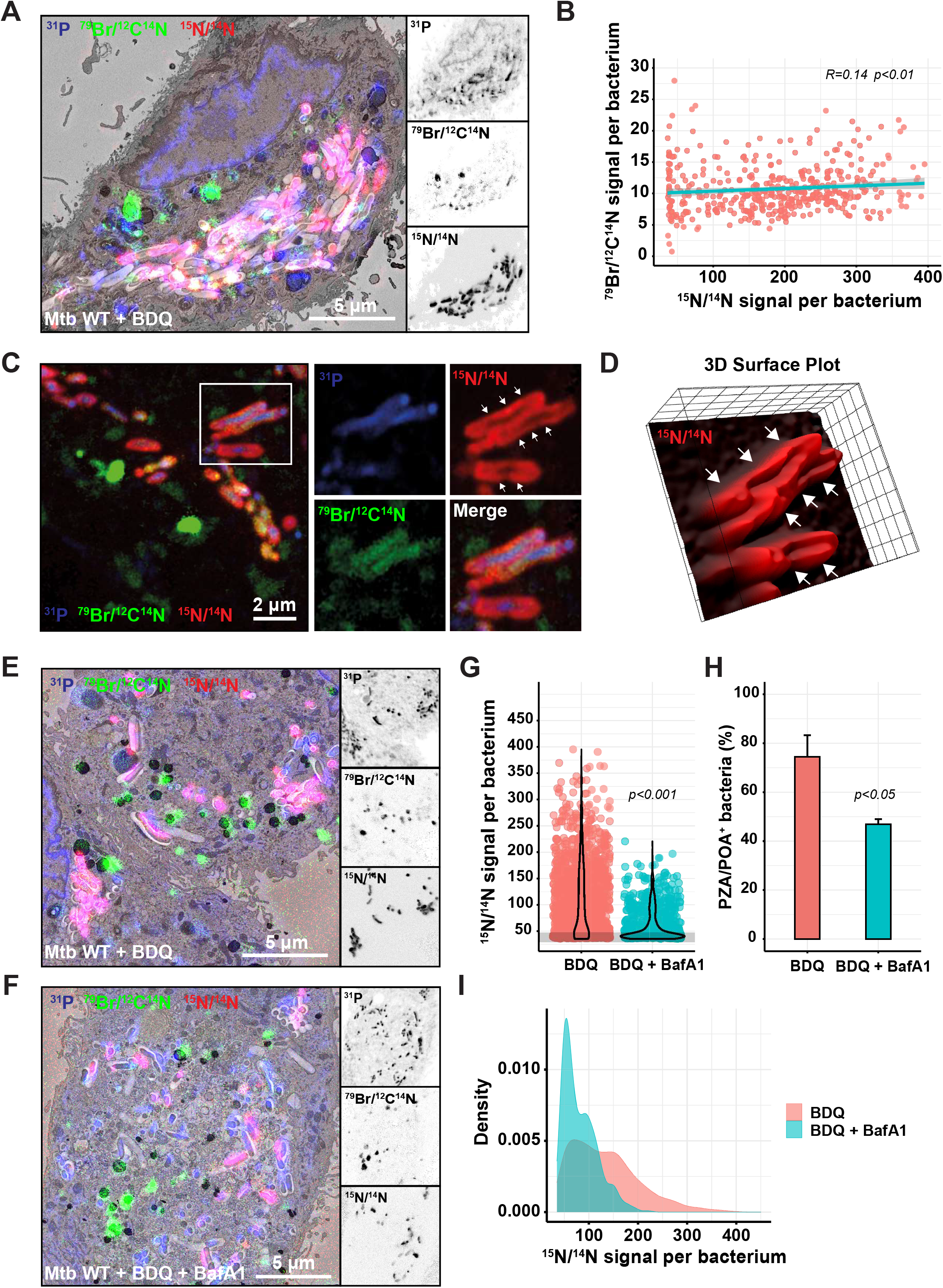
Dual subcellular antibiotic visualisation shows that BDQ enhances PZA accumulation within Mtb-infected MDM. **(A)** Mtb-infected MDM treated with 30 mg/L [^15^N_2_, ^13^C_2_]-PZA and 2.5 mg/L of BDQ for 24 hours. EM micrograph is overlaid with ^31^P (blue), ^79^Br/^12^C^14^N (green) and ^15^N/^14^N (red) NanoSIMS images. Magnifications show ^31^P (top panel), ^79^Br/^12^C^14^N (middle panel) and ^15^N/^14^N (bottom panel) individual signals at the single bacterial-cell level. Scale bar corresponds to 5 μm. **(B)** Pearson’s correlation between ^15^N/^14^N (x-axis) and ^79^Br/^12^C^14^N (y-axis) signals in individual bacterium. Results are from 453 intracellular Mtb WT obtained from 2 biological replicates. Correlation coefficient (R) shown as the cyan line was determined as 0.14 and the *p*-value was calculated from the linear model. **(C)** NanoSIMS images of Mtb-infected MDM treated with 30 mg/L [^15^N_2_, ^13^C_2_]-PZA and 2.5 mg/L of BDQ for 24 hours. Images were filtered with a median filter using the OpenMIMS plugin in FIJI and pseudo-colours are displayed as in **(A)**. Magnifications show the region of interest highlighted by white rectangle. White arrows indicate the peripheral signal of PZA/POA **(D)** 3D surface plot of the ^15^N/^14^N signal from the bacteria depicted in (C) was generated using the 3D surface plot from FIJI. White arrows show the PZA/POA accumulation in 3D. **(E-F)** Mtb-infected MDM treated with 30 mg/L [^15^N_2_, ^13^C_2_]-PZA and 2.5 mg/L BDQ in the presence or absence of 100 nM BafA1 for 24 hours. EM micrograph and magnifications are displayed as in **(A)**. **(G)** Quantitative analysis of ^15^N/^14^N signal per bacterium treated with BDQ alone or in combination with 100 nM BafA1 shown as violin plot with single dots. Grey line indicates the natural background level of the ^15^N/^14^N enrichment. Results were obtained from 810-1275 individually segmented bacteria and p-values were calculated from the linear model. **(H)** Quantification of the percentage of PZA/POA positive (PZA/POA^+^) bacteria. Results are expressed as mean ± standard error of the mean from 2-3 biological replicates and p-values were calculated from the linear model. **(I)** Analysis of the ^15^N/^14^N signal profile of Mtb WT PZA/POA positive bacteria treated with BDQ alone or in combination with 100nM BafA1.

During PZA-BDQ co-treatment, we also observed that high intrabacterial PZA/POA levels had a particular spatial distribution within intracellular Mtb. High-resolution ion micrographs of ^31^P, ^79^Br/^12^C^14^N and ^14^N/^15^N signal analysis and 3D surface plot reconstruction showed that PZA/POA signal was mainly detected at the periphery of the bacterial cell suggesting a strong association with the bacterial cell wall (**Figure 5C-D**). In contrast, the ^79^Br/^12^C^14^N signal corresponding to the BDQ signal was diffuse through the bacterial cytosol (**Figure 5C**).

We then tested if the effect of BDQ on PZA accumulation is reversed by inhibition of endolysosomal acidification. Human macrophages were infected with Mtb WT and incubated with labelled PZA and BDQ in the presence or absence of 100 nM BafA1. The inhibition of endolysosomal acidification decreased PZA/POA levels associated with Mtb (**Figure 5E-G**) (average mean intensity per positive bacteria of 138.84 ± 2.27 *vs* 82.19 ± 1.75, *p*<0.001). Moreover, the percentage of positive bacteria upon BafA1 treatment decreased approximately 1.6-fold (**Figure 5H**). Confirming these observations, distribution of the ^15^N/^14^N enrichment of the population significantly changed after BafA1 treatment (**Figure 5I**). Altogether, by simultaneously imaging two antibiotics, we show that BDQ enhances PZA accumulation within Mtb through endolysosomal targeting, and that the synergistic effect is partially counteracted by inhibiting the activity of vacuolar-type H^+^-ATPase.

## Discussion

By using correlative imaging of antibiotics at the subcellular level, here we provide evidence supporting the idea that distinct antibiotics will differentially accumulate in intracellular compartments with direct consequences for efficacy. The frontline anti-TB drug PZA distributed very differently from BDQ, which was shown to accumulate in host cell lipid droplets ^38^. PZA/POA primarily localises in intracellular Mtb and not in any other organelles or the cytosol, though we cannot rule out that some PZA/POA was washed out during sample processing. PZA/POA was only detectable by the enrichment of ^15^N relative to ^14^N in treated samples but not by the ^13^C label, likely due to matrix effects. The ^15^N isotopic label used during PZA synthesis was positioned in the pyrazine ring and did not allow discriminating between PZA and POA in this study. Being able to distinguish the prodrug from its active form will be important to further define PZA mode of action in intracellular Mtb.

Given that PZA is an effective antitubercular drug *in vivo,* we expected that it would homogenously accumulate in Mtb-infected macrophages at drug concentrations close to the *C*_max_. However, we observed that not all macrophages had detectable levels of PZA/POA and some macrophages were completely devoid of the antibiotic. The possibility that sample processing can differentially affect macrophages in the same population is very low. In this context, very little is known about the mechanisms that allow internalisation of antibiotics in infected cells ^48,49^ and we postulate that the phenotypic state of individual cells determines drug uptake and enrichment.

There was also a highly heterogenous distribution of Mtb-associated PZA/POA signal, strongly suggesting that at physiological levels, PZA does not accumulate in all intracellular Mtb. The observation that in necrotic cells, where permeability of host membranes is higher, very few bacteria were enriched with PZA/POA agrees with *in vivo* studies ^23,50,51^. Although PZA/POA distributes across necrotic tissue, the biophysical parameters of these specific lesions, such as neutral pH, has been proposed to be a key factor linked to PZA/POA inefficacy against Mtb in mouse and guinea pig models ^50,51^.

Intracellular Mtb that is mostly cytosolic at the time of treatment is partially protected from PZA action, suggesting that the neutral pH of cytosol could be detrimental for PZA efficacy. On the other hand, membrane bound localisation renders Mtb more susceptible to PZA. It is likely that ESX-1-mediated phagosomal membrane damage affects the pH surrounding Mtb ^41^. Indeed, Mtb harbouring mutations in the ESX-1-encoding genes fail to prevent phagosome acidification. Furthermore, Mtb mutants in the *phoPR* two-component system, which blocks the secretion of EsxA ^52,53^, were localised in acidic phagosomes. Pharmacological inhibition of the ESX-1 secretion system results in more Mtb localising in acidic compartments ^42^. Thus, ESX-1-mediated secretion reduces exposure of Mtb to acidic compartments and might explain the higher accumulation of PZA/POA within intracellular Mtb ΔRD1 compared to cytosolic Mtb WT. We cannot exclude that other factors potentially different between Mtb WT and Mtb ΔRD1 might be responsible for these differences. Given that side-by-side comparison experiments *in vitro* did not show major changes in susceptibility between Mtb WT and Mtb ΔRD1, it is likely that the host cell microenvironment is the main driver of the ESX-1 dependent alterations in PZA accumulation.

Phagosome acidification is also required for PZA/POA accumulation in intracellular Mtb, suggesting that this antibiotic acts by a host cell-driven pH-dependent mechanism. Changes in pH or proteolytic activity can modulate drug efficacy in mouse macrophages ^14^ and our results indicate that acidification and proteolytic activity contributes to Mtb restriction in human macrophages as well. This agrees with data showing that inhibition of endolysosomal acidification increases mycobacterial replication ^45,54^.

Importantly, both Mtb ESX-1-dependent localisation and phagosome acidification are required for efficient PZA efficacy, as shown by intracellular antibiotic susceptibility assays. PZA efficacy was significantly reduced with inhibitors of acidification, and this effect was more pronounced in Mtb WT than in Mtb ΔRD1. These differences might be explained by the ability of the Mtb WT to efficiently localise in the neutral pH of the cytosol, arguing that cytosolic access of Mtb in human macrophages results in lower efficacy of PZA. Strikingly, in conditions where acidification was impaired, PZA was still able to restrict Mtb WT and Mtb ΔRD1 replication in a dose dependent manner, arguing that PZA/POA also targeted bacteria in non-acidic environments. In fact, compelling evidence has highlighted the efficacy of PZA in non-acidic conditions ^55,56^.

By using a correlative approach that enabled imaging of both PZA and host acidic microenvironments, we discovered that PZA can accumulate in bacteria contained in non-acidic compartments. However, the dynamics of Mtb infection in human macrophages is complex ^40,57^ and it will be important to define at which stage the accumulation of the antibiotic happens with higher temporal resolution. Further studies will define whether time of residence in specific microenvironments define drug enrichment and efficacy. Other bacterial factors that are only expressed within host cells can potentially affect PZA accumulation in acidic and non-acidic environments need to be considered. For example, intracellular bacterial fitness, metabolism and drug influx/efflux activity might influence antibiotic accumulation and efficacy. Altogether, our data show that the heterogeneity of PZA enrichment in Mtb is likely the result of a combination of both bacterial and host cell factors.

BDQ, an antibiotic that targets the mycobacterial ATP synthase and also displays antibacterial activity by increasing lysosomal activity, significantly increased PZA/POA accumulation in Mtb. We did not observe any correlation between the accumulation of BDQ and PZA/POA per single bacterium suggesting that the two drugs may have a cooperative mechanism of action as previously suggested ^23^. Moreover, there was no synergistic effect *in vitro* either at neutral or acidic pH, arguing that the effect of BDQ is host-mediated as previously shown ^47^. Both our *in vitro* and *in cellulo* experiments suggest that BDQ-induced reduction of ATP levels, does not impair active influx/efflux of PZA/POA, however further experiments are required to confirm these findings. Accordingly, inhibition of acidification significantly reduced PZA/POA levels and the proportion of PZA/POA positive bacteria in infected cells suggesting that the observed host-mediated mechanism triggered by BDQ is pH-driven.

The simultaneous subcellular imaging of two structurally unrelated antibiotics allowed us to define intrabacterial distribution of the drugs. Unexpectedly, we found that PZA/POA spatial distribution within individual bacteria was primarily associated with the periphery of the bacillus. Although the biological significance is unclear, it is tempting to postulate that such accumulation is linked to the biological target(s) of the PZA/POA. Protonated POA is able to accumulate and further disrupt membrane potential, membrane transport and intrabacterial pH ^31^. Recent structural and biochemical studies, clearly demonstrated that POA molecules are also able to covalently bind the Mtb protein PanD *in vitro* ^35,58^. Although the bacterial localisation of PanD is unknown, it is possible that CoA biosynthesis enzyme might localise at the inner leaflet of the bacterial cytoplasmic membrane to meet the requirement of specific lipid biosynthesis pathways as it was previously shown between the FAS-II and the mycolic acid synthesis machineries ^59^. Thus, host cell environments and Mtb intracellular localisation affect antibiotic efficacy, arguing that different antibiotics are needed to target heterogeneous intracellular bacterial populations. These results provide an explanation for the observed sterilising activity of PZA in the clinic but also contributes to a conceptual framework for a host cell dependent combined drug therapy in the context of TB. It is likely that many other antibiotics used for the treatment of intracellular pathogens display localisation-dependent potency and efficacy.

## Material and Methods

### Bacterial strains and growth conditions

*M. tuberculosis* H37Rv pTEC19 (Mtb WT) and *M. tuberculosis* H37Rv ΔRD1 pTEC19 (Mtb ΔRD1) strains were used in this study ^40^. Recombinant strains harbouring pTEC19 plasmid (Addgene, #30178) and producing the fluorescent protein E2-Crimson were grown in Middlebrook 7H9 broth supplemented with 0.2% glycerol (v/v) (Fisher Chemical, G/0650/17), 0.05% Tween-80 (v/v) (Sigma-Aldrich, P1754) and 10% ADC (v/v) (BD Biosciences, 212352). Both strains were previously tested for PDIM positivity by thin layer chromatography of lipid extracts from Mtb cultures ^40^. Hygromycin B (Invitrogen, 10687010) was used at a concentration of 50 mg/L as a selection marker for the fluorescent strains. Bacterial cultures (10 mL) were incubated at 37°C with rotation in 50 mL conical tubes.

### Antibiotic susceptibility testing in broth

Susceptibility testing of Mtb WT and Mtb ΔRD1 was performed by end-point determination of optical density at 600 nm (OD_600nm_) using the broth microdilution method in 96-well clear flat-bottom microplates (Corning, 353072). Both Mtb WT and Mtb ΔRD1 strains were grown in complete Middlebrook 7H9 broth until mid-exponential phase (OD_600nm_ = 0.6 ± 0.2) and further adjusted to a bacterial density of approximately 5 × 10^6^ bacteria/mL (OD_600nm_ = 0.05) assuming that an OD_600nm_ of 1 approximates to 10^8^ bacteria/mL. A volume of 100 μL of inoculum was then added to each well containing 100 μL of two-fold serial dilutions of PZA (Sigma, P7136) ranging from 4 to 512 mg/L in complete Middlebrook 7H9 broth, thus giving a final bacterial load of approximately 2.5 × 10^6^ bacteria/mL (OD_600nm_ = 0.025). In addition, wells containing 200 μL of 7H9 medium only were used as sterility/background controls, whereas wells containing 5 mg/L or 2.5 mg/L of RIF (LKT laboratories, R3220) and BDQ (MedChemExpress, HY-14881) respectively, were used as positive controls for growth inhibition. Plates were sealed in zipped-lock bags and incubated at 37°C without agitation. After 14 days, plates were scanned and OD_600nm_ was determined using a microplate reader (Berthold, Mithras^2^ LB 943). Results were expressed as relative growth where Mtb WT and Mtb ΔRD1 growth in complete Middlebrook 7H9 broth was considered as 100%. Experiments were performed in biological duplicate with at least 6 technical replicates.

### Preparation of human-monocyte derived macrophages

Human monocyte-derived macrophages (MDM) were prepared from Leukocyte cones (NC24) supplied by the NHS Blood and Transplant service as previously described ^10,38^. White blood cells were isolated by centrifugation on Ficoll-Paque Premium (GE Healthcare, 17-5442-03) for 60 min at 300 *g.* Mononuclear cells were collected and washed twice with MACS rinsing solution (Miltenyi, 130-091-222) to remove platelets and red blood cells. The remaining were lysed by incubation with 10 mL RBC lysing buffer (Sigma, R7757) per pellet for 10 min at room temperature. Cells were washed with rinsing buffer and then were re-suspended in 80 μL MACS rinsing solution supplemented with 1% BSA (Miltenyi, 130-091-376) (MACS/BSA) and 20 μL anti-CD14 magnetic beads (Miltenyi, 130-050-201) per 10^8^ cells. After 20 min on ice, cells were washed in MACS/BSA solution and re-suspended at a concentration of 10^8^ cells/500 μL in MACS/BSA solution and further passed through an LS column (Miltenyi, 130-042-401) in the field of a QuadroMACS separator magnet (Miltenyi, 130-090-976). The LS column was washed three times with MACS/BSA solution, then CD14 positive cells were eluted, centrifuged and re-suspended in complete RPMI 1640 with GlutaMAX and HEPES (Gibco, 72400-02), 10% foetal bovine serum (Sigma, F7524) and 10 ng/ml of hGM-CSF (Miltenyi, 130-093-867). Cells were plated at a concentration of 10^6^ cells/mL in untreated petri dishes. Dishes were incubated in a humidified 37°C incubator with 5% CO2. After three days, an equal volume of fresh complete media including hGM-CSF was added. Six days after the initial isolation, differentiated macrophages were detached in 0.5 mM EDTA in ice-cold PBS using cell scrapers (Sarsted, 83.1830), pelleted by centrifugation and resuspended in RPMI medium containing 10% foetal bovine serum where cell count and viability was estimated (BioRad, TC20™ Automated Cell Counter) before plating for experiments.

### Macrophage infection with Mtb

For macrophage infection, Mtb WT and Mtb ΔRD1 inoculum were prepared as previously described ^10,38^. Approximately 10 mL of mid-exponential phase bacterial cultures (OD_600nm_ = 0.6 ± 0.2) were centrifuged and washed twice in sterile PBS buffer (pH 7.4). Then, an equivalent volume of sterile 2.5-3.5 mm autoclaved glass beads was added to the pellet and bacterial clumps were disrupted by vigorously shaking. Bacteria were resuspended in cell culture media and the remaining clumps were removed by slow-speed centrifugation at 300 *g* for 5 min. The supernatant was transferred to a fresh tube and OD_600nm_ measured to determine bacterial concentration, assuming once again that an OD_600nm_ of 1 approximates to 10^8^ bacteria/mL. For intracellular antibiotic assays, macrophages were infected with Mtb WT and Mtb ΔRD1 at a multiplicity of infection (MOI) of 1 for 2 h at 37°C. For ion microscopy and full-correlative experiments an MOI of 5-10 was used. After 2 h of uptake, cells were washed twice with PBS to remove extracellular bacteria and fresh media was added.

### Intracellular antibiotic activity assays in Mtb-infected macrophages

Intracellular antibiotic activity assays were performed as previously reported ^38^. Briefly, 3.5 × 10^4^ cells per well were seeded into an olefin-bottomed 96-well plate (Perkin Elmer, 6055302) 16-20 h prior to infection. Cells were infected as described above for 24 h and the culture media was replaced by fresh media containing increasing concentrations of PZA, RIF, INH or left untreated. When indicated, fresh medium containing 100 nM Bafilomycin-A1 (BafA1) (Enzo Life Sciences, BML-CM110-0100) or 100 nM Concanamycin A (ConA) (Sigma-Aldrich, C9705) was added together with the antibiotics. At the required time points, infected cells were washed twice with PBS buffer (pH 7.4) and fixed with a 4% methanol-free paraformaldehyde (Electron Microscopy Sciences, 15710) in PBS buffer (pH 7.4) for 16-20 h at 4°C. Fixative was removed and cells were washed two times in PBS buffer (pH 7.4) before performing DAPI (Invitrogen, D1306) staining for nuclear visualisation. Image acquisition was performed with the OPERA Phenix high-content microscope with a 40x water-immersion 1.1 NA objective. The confocal mode with default autofocus and a binning of 1 was used to image multiple fields of view (323 μm × 323 μm) from each individual well with 10% overlapping, where acquisition was performed at 4 distinct focal planes spaced with 1 or 2 μm. DAPI stained nuclei were detected using λ_ex_ = 405 nm/λ_em_ = 450 nm, where the laser power was set at 20% with an exposure time of 200 ms. E2-Crimson bacteria were detected using λ_ex_ = 595 nm/λ_em_ = 633 nm, the laser power was set at 30% with an exposure time of 200 ms. Analysis was performed using the Harmony software (Perkin Elmer, version 4.9) where maximum projection of the 4 z-planes was used to perform single cell segmentation by using the “Find nuclei” and “Find cells” building blocks. Cells on the edges were excluded from the analysis. The E2-Crimson signal was detected using the “Find Image Region” building block where a manual threshold was applied to accurately perform bacterial segmentation. The Mtb area per cell was determined by quantifying the total area (expressed in μm^2^) of E2-Crimson^+^ signal per single macrophage. The relative growth inhibition was determined by using the following formula ~ (Mean Mtb area per cell t96h - Mean Mtb area per cell t24h) / (Mean Mtb area per cell t24h) and the relative values were obtained by using the untreated control as a reference of 100% growth (0% inhibition). All the results were exported as CSV files, imported in the R studio software (The R Project for Statistical Computing) and graphs were plotted with the ggplot2 package.

### Resin embedding and block trimming

Mtb-infected cells were washed once in HEPES 0.2 M buffer, fixed with 2.5% glutaraldehyde (Sigma, G5882), 4% methanol-free paraformaldehyde (Electron Microscopy Sciences, 15710) in 0.2 M HEPES (pH 7.4) for 16-20 h at 4°C and further processed for Scanning Electron Microscopy (SEM) and nanoscale secondary ion mass spectrometry (NanoSIMS) in a Biowave Pro (Pelco, USA) with use of microwave energy and vacuum. Cells were washed twice in HEPES 0.2 M buffer (Sigma-Aldrich, H0887) at 250 W for 40 s, post-fixed using a mixture of 2% osmium tetroxide (Taab, O011) and 1.5% potassium ferricyanide (Taab, P018) (v/v) at equal ratio for 14 min with power set up at 100 W (with/without vacuum 20” Hg at 2-min intervals). Samples were washed with distilled water twice on the bench and twice again in the Biowave 250 W for 40 s and further stained in 1% thiocarbohydrazide (Sigma-Aldrich, 223220) in distilled water (w/v) for 14 min as described above. After repeated washes, samples were also stained with 2% osmium tetroxide (Taab, O011) in distilled water (w/v) for 14 min with similar settings. Samples were washed 4 times as before and stained with 1% aqueous uranyl acetate (Agar scientific, AGR1260A) in distilled water (w/v) for 14 min and then washed again.

Samples were dehydrated using a step-wise ethanol series of 50, 75, 90 and 100%, then washed four times in absolute acetone at 250 W for 40 s per step. Samples were infiltrated with a dilution series of 25, 50, 75 and 100% Durcupan ACM® (Sigma-Aldrich, 44610) (v/v) resin to acetone. Each step was for 3 min at 250 W power (with/without vacuum 20” Hg at 30 s intervals). Samples were then cured for a minimum of 48 h at 60°C. The sample block was trimmed, coarsely by a razor blade then finely trimmed using a 35° ultrasonic, oscillating diamond knife (DiATOME, Switzerland) set at a cutting speed of 0.6 mm/s, a frequency set by automatic mode and a voltage of 6.0 V, on an ultramicrotome EM UC7 (Leica Microsystems, Germany) to remove all excess resin surrounding an area for analysis.

For bacterial pellets, fixed bacteria were pelleted at 3000 *g* for 5 min and washed twice in HEPES 0.2 M buffer (Sigma-Aldrich, H0887) at 250 W for 40 s, post-fixed using a mixture of 2% osmium tetroxide (Taab, O011) and 1.5% potassium ferricyanide (Taab, P018) (v/v) at equal ratio for 14 min with power set up at 100 W (with/without vacuum 20” Hg at 2 min intervals). Samples were washed twice with distilled water in the Biowave at 250 W for 40 s and further stained with 1% aqueous uranyl acetate (Agar scientific, AGR1260A) in distilled water (w/v) for 14 min and then washed again as before. Between steps, bacteria were centrifuged at 3000 *g* for 5 min. Samples were dehydrated using a stepwise ethanol series of 50, 75, 90 and 100%, then washed four times in absolute acetone at 250 W for 40 s per step. Samples were infiltrated with a dilution series of 25, 50, 75 and 100% Durcupan ACM® (Sigma-Aldrich, 44610) (v/v) resin to acetone. Each step was for 3 min at 250 W.

### Nanoscale secondary ion mass spectrometry (NanoSIMS)

For ion microscopy experiments, cells were infected for 24 h and the culture media was replaced by fresh media containing either 30 μg/mL [^15^N_2_, ^13^C_2_]-PZA alone or in combination with 2.5 μg/mL BDQ or 100 nM BafA1 for an additional 24 h. Samples were fixed, embedded in resin and trimmed as described above before performing acquisition. The sections were imaged by SEM and NanoSIMS as previously described ^38^. Briefly, 500 nm sections were cut using ultramicrotome EM UC7 (Leica Microsystems, Germany) and mounted on 7 mm × 7 mm silicon wafers. Sections on silicon wafers were imaged using an FEI Verios SEM (Thermo Fisher Scientific, USA) with a 1 kV beam with the current at 200 pA. The same sections were then coated with 5 nm gold and transferred to a NanoSIMS 50 or 50L instrument (CAMECA, France). The regions that were imaged by SEM were identified using the optical microscope in the NanoSIMS. A focused ^133^Cs+ beam was used as the primary ion beam to bombard the sample; secondary ions (^12^C^−^, ^12^C^14^N^−^,^12^C^15^N^−^,^31^P^−^ and ^79^Br^−^) and secondary electrons were collected. A high primary beam current of ~1.2 nA was used to scan the sections to remove the gold coating and implant ^133^Cs^+^ to reach a dose of 1×10^17^ ions/cm^2^ at the steady state of secondary ions collected. Identified regions of interest were imaged with a ~3.5 pA beam current and a total dwell time of 10 ms/pixel. Scans of 512 × 512 pixels were obtained. Quantification of secondary ion signal intensities was performed using the open source software ImageJ/Fiji v1.53a and the OpenMIMS v3.0.5 plugin. Briefly, bacterial detection and manual segmentation was performed based on the ion signal from ^31^P and corresponding correlative EM images to identify single bacterial cells. Quantification of intrabacterial PZA/POA levels within region of interests was performed using ^15^N/^14^N ratio calculated from ^12^C^15^N/^12^C^14^N signals, where the signal from ratiometric images (512 × 512 pixels, 32-bit) defined the ion level of individual bacteria (expressed as parts per 10^4^). The ^15^N/^14^N ratio value presented is multiplied by 10,000. Background determination of natural ^15^N levels was performed by quantifying ^15^N/^14^N ratio within untreated or DMSO-treated samples. The obtained control values were tested for normal distribution by performing a quantile-quantile plot followed by a Shapiro-Wilk test and the mean background value ± standard deviation was determined. The statistical empirical rule was used and the mean background value ± 3*standard deviation (μ ± 3*σ) was applied to define a background level with a 99.7% confidence. All mean ^15^N/^14^N ratiometric values obtained above this threshold were subsequently considered as PZA/POA positive bacteria. A similar strategy was used to quantify intrabacterial BDQ levels except that detection and quantification was performed using normalized signal from ^79^Br/^12^C^14^N. Ion values from individual bacteria in each experimental condition were exported as CSV files, imported in R (The R Project for Statistical Computing) and graphs were plotted with the ggplot2 package

### Correlative Light Electron Ion Microscopy (CLEIM)

Mtb-infected macrophages were treated with 30 μg/mL [^15^N_2_, ^13^C_2_]-PZA for 24 h, then washed once with media and stained with 200 nM LysoTracker™ Red DND-99 (Invitrogen, L7528) for 30 min in a humidified 37°C incubator with 5% CO2. Subsequently, cells were washed twice with 0.2 M HEPES buffer (pH 7.4) and fixed in 0.1% glutaraldehyde (Sigma, G5882), 4% methanol-free paraformaldehyde (Electron Microscopy Sciences, 15710) in 0.2 M HEPES (pH 7.4) for 16-20 h at 4°C. Fixative was removed and cells were washed three times in 0.2 M HEPES (pH 7.4) before performing DAPI staining for nuclear visualisation. Image acquisition was performed using Leica SP8 confocal microscope equipped with a HC PL APO CS2 63x/1.40 oil objective. Images of 1024 × 1024 pixels were acquired with Diode 405 nm, DPSS 561 nm and HeNe 633 nm lasers where intensities were set up as 2.5%, 5% and 6% respectively. Emitted signal from each channel was collected at 450 +/− 30 nm, 585 +/− 15 nm and 710 +/-15 nm for DAPI, LysoTracker and E2-Crimson respectively. The cells of interest were then resin embedded, the block was trimmed, SEM/NanoSIMS acquisition and analysis was performed as described above. For correlation, fluorescent images, EM and nanoSIMS micrographs were converted to tiff files and linear adjustments made to brightness and contrast using ImageJ/Fiji v1.53a. Images were further aligned to EM micrographs with Icy 2.0.3.0 software (Institut Pasteur, France), using the ec-CLEM Version 1.0.1.5 plugin. No fewer than 10 independent fiducials were chosen per alignment for 2D image registration. When the fiducial registration error was greater than the predicted registration error, a non-rigid transformation (a nonlinear transformation based on spline interpolation, after an initial rigid transformation) was applied as previously described ^60^. Images were finally displayed using the open source software ImageJ/Fiji v1.53a.

### Cytotoxicity and endolysosomal proteolytic assays

Approximately 3.5 × 10^4^ MDM per well were seeded into an olefin-bottomed 96-well plate (Perkin Elmer, 6055302) in complete RPMI medium. After 16-20 h, the medium was replaced by fresh medium containing 100 nM Bafilomycin-A1 for 24-72 h. To evaluate cytotoxicity, cells were stained for 30 min using the Blue/Green ReadyProbes™ Cell Viability Imaging Kit (Invitrogen, R37609) following the manufacturer recommendations. A positive control was done by adding 50 mM hydrogen peroxide (Sigma, 18304) to the staining solution and live-cell imaging was further performed using the OPERA Phenix microscope with a 40x water-immersion objective. Segmentation and analysis were performed using the Harmony software (Perkin Elmer, version 4.9) where maximum projection of 4 individual z-planes with an approximate distance of 2 μm was used to perform single cell segmentation by combining the “Find nuclei” and “Find cells” building blocks. Cell viability was determined by counting the number of nuclei in each experimental condition and the proportion of green positive nuclei, where a threshold of 5 × 10^3^ fluorescent arbitrary units was applied to define positive cells. For endolysosomal proteolytic activity assays, cells were pulsed with 10 μg/ml of DQ™ Red BSA (Invitrogen, D12051) for 4 h as previously described ^61^. Stained cells were then washed twice and reincubated in complete RPMI medium containing 1 μg/ml Hoechst 33342 (Immunochemistry Technologies, 639) and live-cell imaging was further performed using the OPERA Phenix microscope with a 40x water-immersion objective. In these assays, all lasers were set at 20 % with an exposure time of 200 ms. After single-cell segmentation, DQ-BSA positive puncta were segmented by using “Find spot” and their respective properties were determined using the “Morphology properties” and “Intensity properties” building blocks. All the results were exported as CSV files, imported in R (The R Project for Statistical Computing) and graphs were plotted with the ggplot2 package

### Live fluorescence imaging and Mtb-LysoTracker co-localisation analysis

For live-cell imaging, cells were infected with Mtb WT and Mtb ΔRD1 at a MOI of 1 as described above. After, 24 h the culture media was replaced by fresh media only or fresh medium containing 100 nM BafA1. Approximately, 24 h after treatment, infected cells were washed once with PBS buffer (pH 7.4) and stained for 30 min with complete RPMI medium containing 200 nM LysoTracker™ Blue DND-22 (Invitrogen, L7525) in a humidified 37°C incubator with 5% CO2. Infected cells were then washed with PBS buffer (pH 7.4) and resuspended in complete RPMI containing NucSpot® Live 488 (Biotium, #40081) following the manufacturer recommendations. Live-cell imaging was further performed using the OPERA Phenix microscope with a 63x water-immersion objective and acquisition settings for blue, green and far-red channels were conserved as described above. Segmentation and analysis were performed using the Harmony software (Perkin Elmer, version 4.9) where the E2-Crimson signal from single z-planes was detected using the “Find Image Region” building block and where a manual threshold was applied to accurately perform bacterial segmentation. Each individual region of interest was transformed into a mask and extended by using the “Find Surrounding Regions” building block with an individual threshold of 0.8 and including the input region. This mask was used to determine the LysoTracker mean intensity associated to the Mtb region. To determine the percentage of LysoTracker positive bacteria, 200 bacterial regions considered as LysoTracker negative were analysed. The control mean intensity values were tested for normal distribution by performing a quantile-quantile plot followed by a Shapiro-Wilk test and the LysoTracker mean intensity value ± standard deviation was determined. The statistical empirical rule was used and the mean background value ± 3*standard deviation (μ ± 3*σ) was applied to define a LysoTracker negative background level with a 99.7% confidence. All mean fluorescence intensity values obtained above this threshold were subsequently considered as LysoTracker positive bacteria.

### Statistical analysis

For NanoSIMS experiments statistical analysis was performed using the linear model as previously described ^38^. Briefly, normalised ion count values were log2 transformed to achieve approximate normality and a (multi-)linear model was applied to predict these log2counts. The following formula was used *“(log2(counts) ~ modulator + donor)”* allowing the consideration of donor-to-donor variability as parameter. Models were fit using the *“lm()”* function in R studio. All *p*-values correspond to the t-statistic assessing significant difference from zero of the model coefficients associated with the reported condition, relative to the baseline control condition. For replication experiments and intracellular antibiotic activity assays, the means between the group of interest were tested for significant differences using one-way ANOVA followed by Tukey’s post-test with the *“aov()”* and *“TukeyHSD()”* functions in R. All the *p*-values contained in the text or the figures are relative to the control condition (unless otherwise stated).

## Acknowledgements

We would like to acknowledge all members of the Host-Pathogen Interactions in Tuberculosis laboratory for insightful discussions. We thank Luiz Pedro S. de Carvalho for reading the manuscript and helpful discussions. We are also grateful to the Advanced Light Microscopy and Electron Microscopy facilities from the Francis Crick Institute for their support in various aspects of this study. This work was supported by the Francis Crick Institute (to MGG), which receives its core funding from Cancer Research UK (FC001092), the UK Medical Research Council (FC001092), and the Wellcome Trust (FC001092). HJ is supported by an Australian Research Council Discovery Early Career Research Award and a Rebecca L Cooper Medical Research Foundation Project Grants. PS has received funding from the European Union’s H2020 research and innovation programme under the Marie Sklodowska-Curie grant agreement *SpaTime_AnTB* n°892859. The authors are also grateful to the Centre for Microscopy, Characterisation & Analysis at the University of Western Australia, which is funded by the University and both the State and Commonwealth Governments.

## Authors contribution

MGG conceived and supervised the project. PS, DJG, HJ and MGG designed the experiments. PS, DJG, AF, KC and HJ performed experiments. All authors analysed data and provided intellectual input. PS and MGG wrote the manuscript with input from DJG, AF and HJ. All authors read the manuscript and provided critical feedback.

## Abbreviations

(Mtb): *Mycobacterium tuberculosis*
(TB): tuberculosis
(INH): isoniazid
(RIF): rifampicin
(ETB): ethambutol
(PZA): pyrazinamide
(BDQ): bedaquiline
(POA): pyrazinoic acid
(CLEIM): correlative light electron and ion microscopy
(OD_600nm_): optical density 600 nm
(MDM): Human monocyte-derived macrophages
(MOI): multiplicity of infection
(SEM): Scanning Electron Microscopy
(NanoSIMS): nanoscale secondary ion mass spectrometry
(*C*_max_): maximum serum concentration
(BafA1): Bafilomycin-A1
(ConA): Concanamycin A.

