## Supplementary information for "Intracellular localisation of *Mycobacterium tuberculosis* affects antibiotic efficacy"


This file contains the Supplementary Figures (S1-S11) and their respective legends

**Supplementary Figures Legend**

**Figure S1: Experimental and analytical electron and ion microscopy workflow**

**(A)** Schematic representation of the experimental procedure followed to perform MDM infection, [$^{15}N_2$, $^{13}C_2$]-PZA treatment, electron/ion microscopy samples processing and images acquisition. **(B)** Schematic representation of the segmentation and analysis pipeline used in this study to perform quantitative analysis of ion microscopy micrographs. The complete procedures are detailed in the *Material and Methods* section of this manuscript.

**Figure S2: Plasma membrane integrity of infected cells contribute to PZA/POA accumulation in Mtb**

**(A)** Representative images of PZA/POA distribution Mtb WT contained within a necrotic cell. Mtb-infected MDM were treated with 30 mg/L [$^{15}N_2$, $^{13}C_2$]-PZA for 24 hours. EM micrograph is overlaid with $^{31}P$ (blue) and $^{15}N/^{14}N$ (red) NanoSIMS images. Magnifications show $^{31}P$ (top panel) and $^{15}N/^{14}N$ (bottom panel) individual images at the single bacterial-cell level. Scale bar corresponds to 10 µm. Region of interest highlighted by the white rectangle, is shown in detail in the right panel. Scale bar corresponds to 2 µm. **(B)** Quantitative analysis of $^{15}N/^{14}N$ signal per bacterium within intact cells (*Cellular*) and necrotic (*Necrotic*) cells, shown as violin plot with single dots. Grey line indicates the background level of the $^{15}N$ enrichment. Results were obtained from 1659 individually segmented bacteria and p-values were calculated from the linear model.

**Figure S3: Intracellular Mtb ΔRD1 localises more often in acidic environments than Mtb WT**

**(A)** Representative fluorescence images of Mtb WT and Mtb ΔRD1-infected MDM stained with LysoTracker in the presence or absence of BafA1. Infected cells were exposed for 24 hours to 100 nM of BafA1 or left untreated, then cells were pulsed with 200 nM of LysoTracker Blue for 30 min and counter stained with NucSpot Green dye. Live cells were imaged live using the OPERA Phenix. Magnifications display nuclear staining (blue), Mtb E2-Crimson (red) and LysoTracker labelling (green). Scale bar corresponds to 20 µm. Region of interests showing Mtb-LysoTracker co-localization are highlighted by the white arrows. **(B)** Quantification of Mtb-associated LysoTracker mean fluorescence intensities. Results are expressed as mean ± standard error of the mean from 2 biological replicates performed at least in three technical replicates. Statistical significance in comparison to the respective control condition or between Mtb WT and Mtb ΔRD1 was assessed by comparing the means of each conditions using one-way ANOVA followed with Tukey's multiple comparisons test. **(C)** Quantitative analysis of LysoTracker mean fluorescence signal per bacterial region of interest shown as violin plot with single dots. The percentage of LysoTracker positive events is displayed on top of the grey boxes. Results were obtained from 2922-10946 individually segmented bacterial region of interest.

**Figure S4: Mtb WT and Mtb ΔRD1 accumulates similar PZA/POA and display identical antibiotic susceptibility towards PZA *in vitro***

**(A)** The activity of PZA onto Mtb WT and Mtb ΔRD1 *in vitro* was tested using the microdilution methods. Susceptibility was assessed using increasing concentration of PZA ranging from 4 to 512 mg/L in complete Middlebrook 7H9 broth, and 5 µg/mL BDQ or 2.5 µg/mL RIF were used as positive inhibition control. After 14 days of incubation, plates were scanned and $OD_{600nm}$ was determined. Results were expressed as relative growth where Mtb WT and Mtb ΔRD1 growth in complete Middlebrook 7H9 broth was considered as 100 %. Grey dashed line indicates 50% of inhibition. Experiments were performed in biological duplicate with at least 6 technical replicates. **(B)** Representative images of PZA/POA accumulation in Mtb WT and Mtb ΔRD1 *in vitro* at pH6.8. Mtb strains were grown until reaching mid-exponentially growing phase in Middlebrook 7H9 medium, inoculated in fresh media adjusted at pH6.8 or pH5.5 and further treated with 30 mg/L [$^{15}N_2$, $^{13}C_2$]-PZA for 24 hours. Micrographs display $^{15}N/^{14}N$ signals as hyperspectral images. Scale bars correspond to 5 µm. Magnifications show $^{31}P$ (top panel) and $^{15}N/^{14}N$ (bottom panel) individual NanoSIMS images. On the right, the quantitative analysis of $^{15}N/^{14}N$ signal per bacterium shown as violin plot with single dots. Grey line indicates the natural background level of the $^{15}N/^{14}N$ enrichment. In that specific context, bacterial segmentation was performed using the $^{31}P$ signal and the "threshold" function from FIJI. The "analyse particles" function from FIJI was used to quantify $^{15}N/^{14}N$ in each region of interest. Results are from 919-1059 individual regions of interests.

**Figure S5: BafA1 treatment impairs endolysosomal proteolytic activity in MDM**

**(A)** Representative fluorescence images of DQ-BSA stained MDM in the presence or absence of BafA1. Approximately 30,000 MDM were exposed for 2, 24 or 72 hours to 100 nM of BafA1 or left untreated, then cells were pulsed with 10 µg/ml of DQ™ Red BSA for 4 hours in the continuous presence of the modulator BafA1 (except the for the control). After 4 hours, cells were washed, stained with Hoechst and imaged live using the OPERA Phenix. Magnifications display nuclear staining (blue) and DQ-BSA labelling (red). Scale bar corresponds to 50 µm. Region of interests highlighted by the white rectangles, are shown in detail in the bottom panels respectively. Scale bar corresponds to 5 µm. **(B)** Quantitative analysis of DQ-BSA mean fluorescence intensity per cell expressed as arbitrary units (AU). **(C)** Quantitative analysis of DQ-BSA spot number normalized per cell area expressed as arbitrary units (AU). Analysis was performed using the “Find spot”, “Morphology properties” and “Intensity properties” building blocks from the Harmony software (Perkin Elmer, version 4.9). From 3420 to 7101 stained MDM were analysed. Results are from 2 biological replicates. Statistical analysis was performed using one-way ANOVA followed by Tukey’s post-test with the “*aov()*” and “*TukeyHSD()*” functions in R. The *p*-values displayed in that figure are all relative to the control condition.

**Figure S6: Inhibition of MDM endolysosomal acidification by BafA1 and ConA doesn’t trigger any cytotoxic effects**

**(A)** Representative fluorescence images of Blue/Green (Live/Dead) stained MDM in the presence or absence of v-ATPase inhibitors for 24 hours. Approximately 30,000 MDM were exposed for 24 hours to 100 nM BafA1, 100 nM ConA or left untreated. Hydrogen peroxide (50 mM) was used as positive control in this assay. Live-cell imaging was performed using the OPERA Phenix microscope with a 40x water-immersion objective. Magnifications display live nuclear staining (blue) and dead nuclear straining (green). Scale bar corresponds to 50 µm. **(B-C)** Quantification of the total number of nuclei detected (y-axis) in each condition after 24 hours. The percentage of green (dead) nuclei in each condition is display on top of each bar chart. Results are expressed as mean ± standard deviation. Each panel represents one biological replicate. **(D)** Representative fluorescence images of Blue/Green (Live/Dead) stained MDM in the presence or absence of v-ATPase inhibitors for

72 hours. Approximately 30,000 MDM were exposed for 72 hours to 100 nM BafA1, 100 nM ConA or left untreated. Magnifications are displayed as described in **(A). (E-F)** Quantification of the total number of nuclei detected (y-axis) in each condition after 72 hours. The percentage of green (dead) nuclei in each condition is display on top of each bar chart. Results are expressed as mean ± standard deviation. Each panel represents one biological replicate.

**Figure S7: Experimental and analytical workflow of high-content intracellular antibiotic susceptibility assays.**

**(A)** Schematic representation of the experimental procedure followed to perform MDM infection, antibiotic treatment, chemical fixation, staining, image acquisition and analysis. **(B)** Schematic representation of the segmentation and analysis pipeline used in this study to perform quantitative analysis of fluorescence microscopy micrographs. The complete procedures are detailed in the *Material and Methods* section of this manuscript.

**Figure S8: Inhibition of MDM endolysosomal acidification by BafA1 and ConA promotes Mtb replication and cell to cell spread.**

**(A and C)** Representative confocal fluorescence images of Mtb WT and Mtb ∆RD1-infected MDM for 24 hours and further treated for 72 hours in the presence or absence of v-ATPase inhibitors. Magnifications display nuclear staining (blue) and Mtb-producing E2-Crimson (red). Scale bar corresponds to 50 µm. **(B and D)** Growth curves of Mtb WT and ∆RD1 strains in Middlebrook 7H9 medium in the presence of 100 nM BafA1, 100 nM ConA or left untreated. Results are representative of two biological replicates. **(E)** Quantitative analysis of E2-Crimson Mtb WT and Mtb ∆RD1 area per single-cell expressed in µm$^2$. Results are displayed in violin plots where grey lines represent the mean Mtb area per cell pre-treatment ($t_{24\ h}$) and black dots represent the mean Mtb area per cell post-treatment ($t_{96\ h}$). From 2124 to 2722 and 3273 to 4775 infected MDM were analysed for Mtb WT and Mtb ∆RD1 respectively. Results are from 2 biological replicates. **(F)** Quantification of the mean proportion of Mtb WT and Mtb ∆RD1 infected cells in the presence or absence of v-ATPase inhibitors 96 h after infection. Results are expressed as mean ± standard deviation. Results are from 2 biological replicates. **(G)** Quantification of the average number of nuclei per field during Mtb WT and Mtb ∆RD1 infection in the presence or absence of v-ATPase inhibitors 96 h after infection. A fixed number of 35 fields were images using the OPERA Phenix microscope with a 40x water-immersion objective. Results are expressed as mean ± standard deviation. Results are from 2 biological replicates.

**Figure S9: PZA inhibition is more effective against Mtb ∆RD1 mutant strain**

**(A-B)** Intracellular replication of Mtb WT and Mtb ∆RD1 in the presence of PZA and 100 nM BafA1 assessed by CFU counting. MDM were infected for 24 hours and further treated for an additional 72 hours with/without 30 mg/L PZA in the presence or absence of v-ATPase inhibitor. Cells were lysed using PBS-Triton X100 0.1%, serially diluted and plated onto 7H11 agar plates. Plates were incubated at 37 °C for 4 weeks. Results are expressed as means of technical triplicate ± standard error of the mean. Representative of two biological replicates.

**Figure S10: Inhibition of MDM endolysosomal acidification by BafA1 doesn't impact Mtb susceptibility towards other anti-TB frontlines drugs.**

**(A-D)** Quantitative analysis of E2-Crimson Mtb WT and Mtb ∆RD1 area per single-cell expressed in $\mu m^2$ in the presence of increasing concentration of RIF (0-5 mg/L) or INH (0-5 mg/L). Results are displayed in violin plots where grey lines represent the mean Mtb area per cell pre-treatment ($t_{24h}$) and black dots represent the mean Mtb area per cell post-treatment ($t_{96h}$). **(A and C)** Antibiotic inhibition assays were performed in the absence of BafA1 whereas 100 nM BafA1 was used in the conditions depicted in **(B and D)**. From 2761 to 4536 and 2487 to 3791 infected MDM were analysed for Mtb WT and Mtb ∆RD1 respectively. Results are from 2 biological replicates.

**Figure S11: BDQ doesn't enhance PZA/POA accumulation *in vitro***

**(A-B)** Representative images of PZA/POA accumulation in Mtb WT *in vitro* in the presence of BDQ. Mtb WT was grown until reaching mid-exponentially growing phase in Middlebrook 7H9 medium, inoculated in fresh media adjusted at pH6.6 or pH5.5 and further treated with 30 mg/L [$^{15}N_2$, $^{13}C_2$]-PZA alone or in combination with 2.5 mg/L BDQ for 24 hours. Micrographs display $^{15}N/^{14}N$ signals as hyperspectral images. Magnifications show $^{31}P$ (top panel) and $^{15}N/^{14}N$ (bottom panel) individual NanoSIMS images. **(C)** Quantitative analysis of $^{15}N/^{14}N$ signal per bacterium shown as violin plot with single dots. Grey line indicates the natural background level of the $^{15}N/^{14}N$ enrichment. In that specific context, bacterial segmentation was performed using the $^{31}P$ signal and the “threshold” function from FIJI. The “analyse particles” function from FIJI was used to quantify $^{15}N/^{14}N$ in each region of interest. Results are from 254-352 individual regions of interests.

### Figure S1

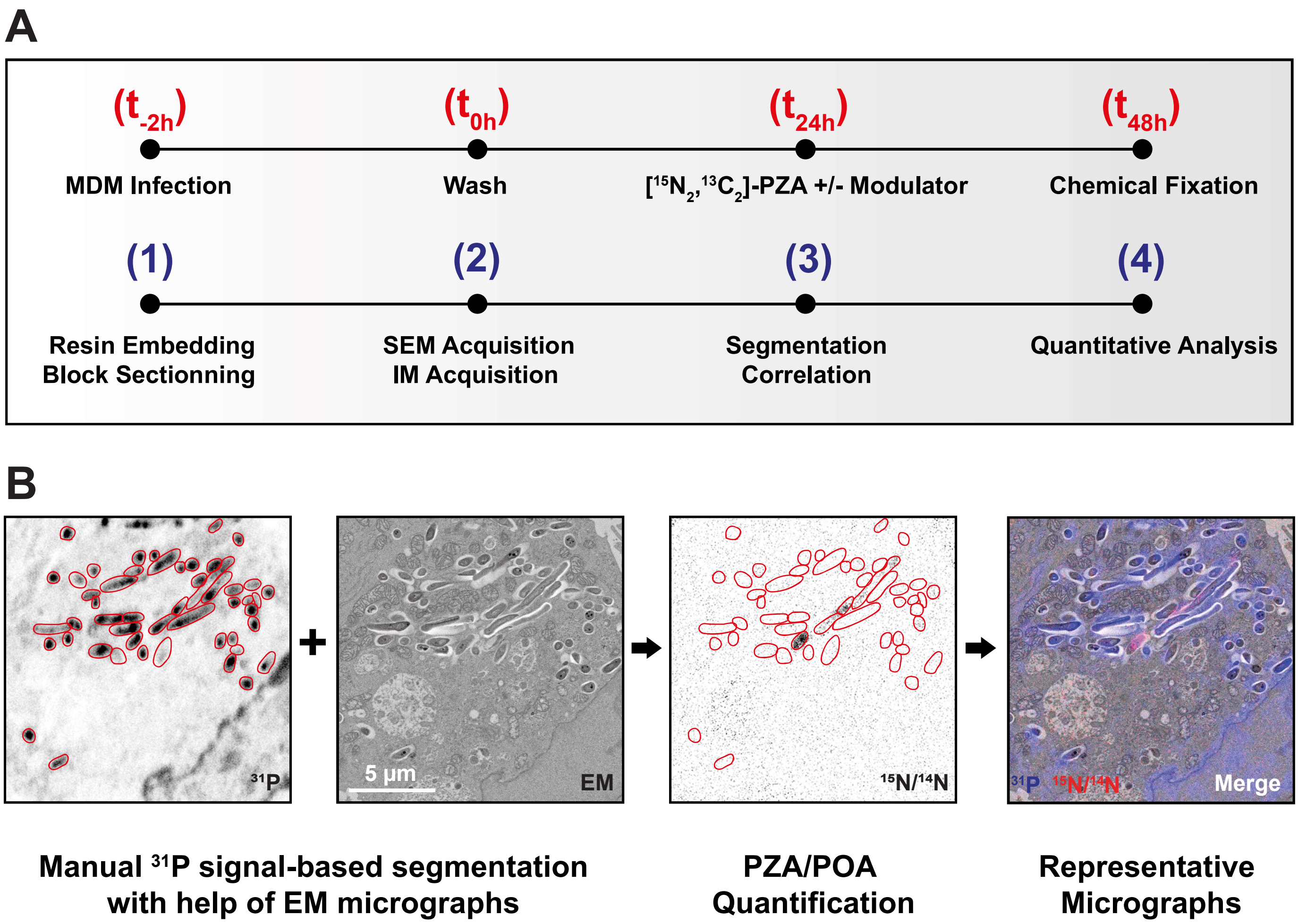
A
(t-2h)
(t0h)
(t24h)
(t48h)
MDM Infection
Wash
[15N2,13C2]-PZA +/- Modulator
Chemical Fixation
(1)
(2)
(3)
(4)
Resin Embedding
Block Sectionning
SEM Acquisition
IM Acquisition
Segmentation
Correlation
Quantitative Analysis
B
31P
5 µm
EM
15N/14N
31P 15N/14N
Merge
Manual 31P signal-based segmentation
with help of EM micrographs
PZA/POA
Quantification
Representative
Micrographs

### Figure S2

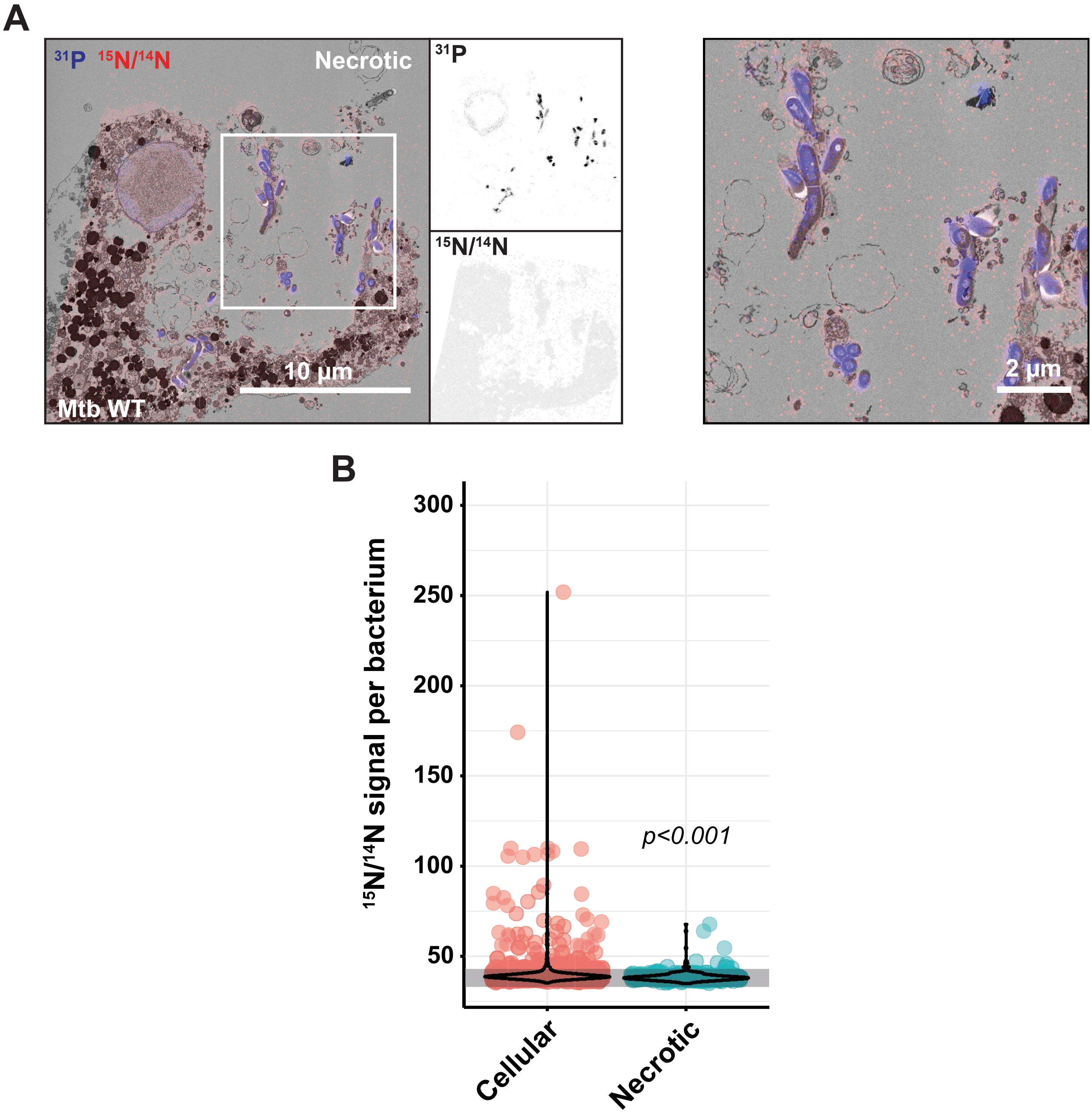

A
31P 15N/14N
Necrotic
10 µm
Mtb WT
31P
15N/14N
2 µm
B
15N/14N signal per bacterium
300
250
200
150
100
50
p<0.001
Cellular
Necrotic

### Figure S3

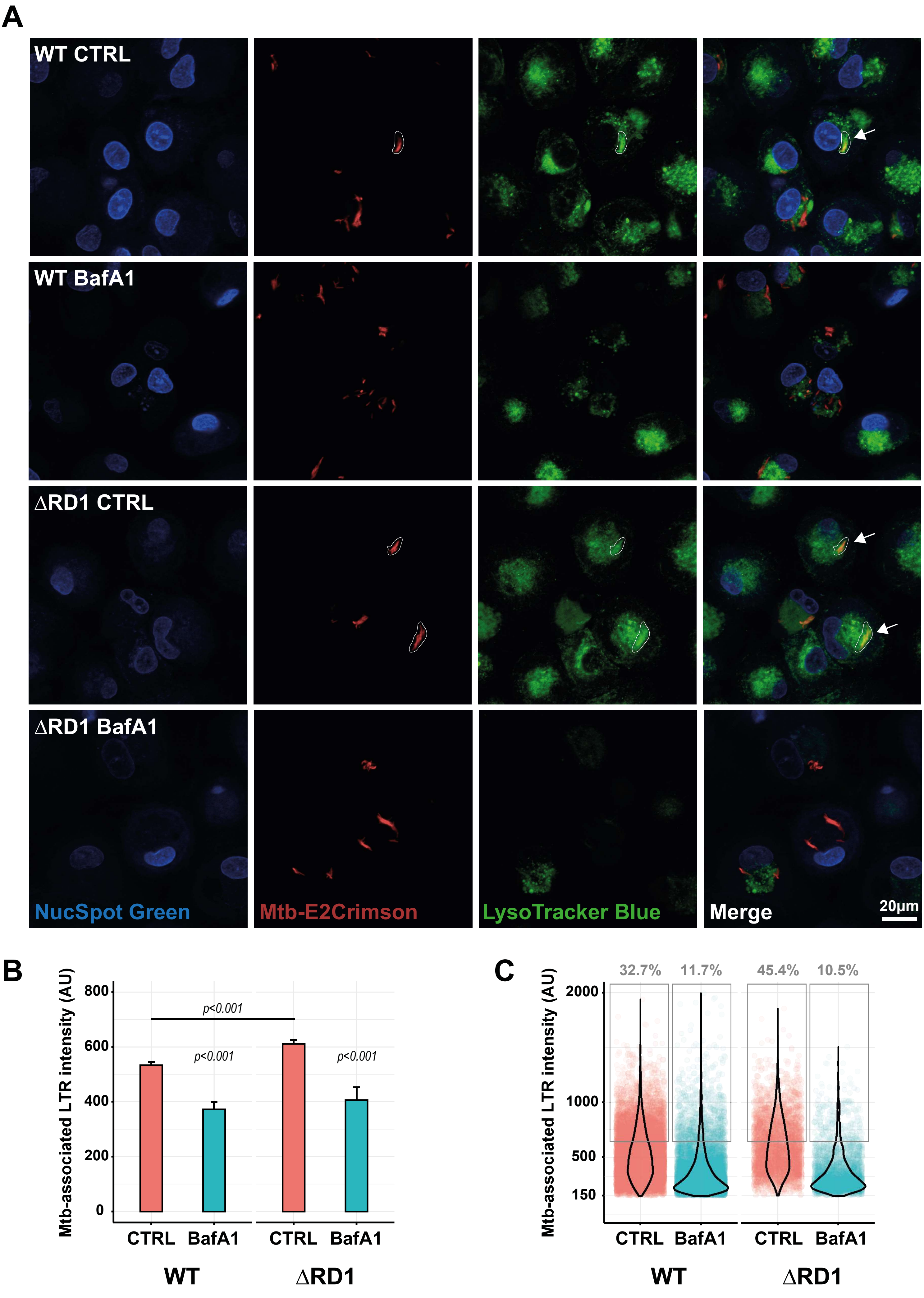
A
WT CTRL
WT BafA1
ΔRD1 CTRL
ΔRD1 BafA1
NucSpot Green
Mtb-E2Crimson
LysoTracker Blue
Merge
20μm
B
Mtb-associated LTR intensity (AU)
800
600
400
200
0
p<0.001
p<0.001
p<0.001
CTRL
BafA1
CTRL
BafA1
WT
ΔRD1
C
Mtb-associated LTR intensity (AU)
2000
1000
500
150
32.7%
11.7%
45.4%
10.5%
CTRL
BafA1
CTRL
BafA1
WT
ΔRD1

### Figure S4

**A**

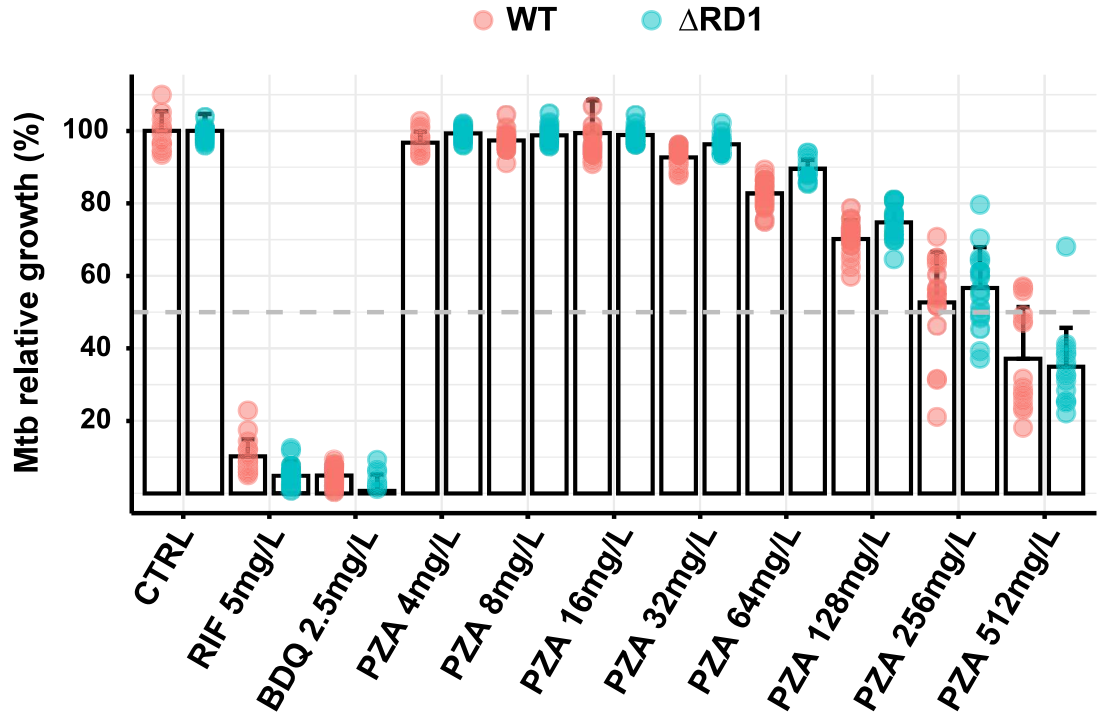
WT
ΔRD1
Mtb relative growth (%)
100
80
60
40
20
CTRL
RIF 5mg/L
BDQ 2.5mg/L
PZA 4mg/L
PZA 8mg/L
PZA 16mg/L
PZA 32mg/L
PZA 64mg/L
PZA 128mg/L
PZA 256mg/L
PZA 512mg/L

**B**

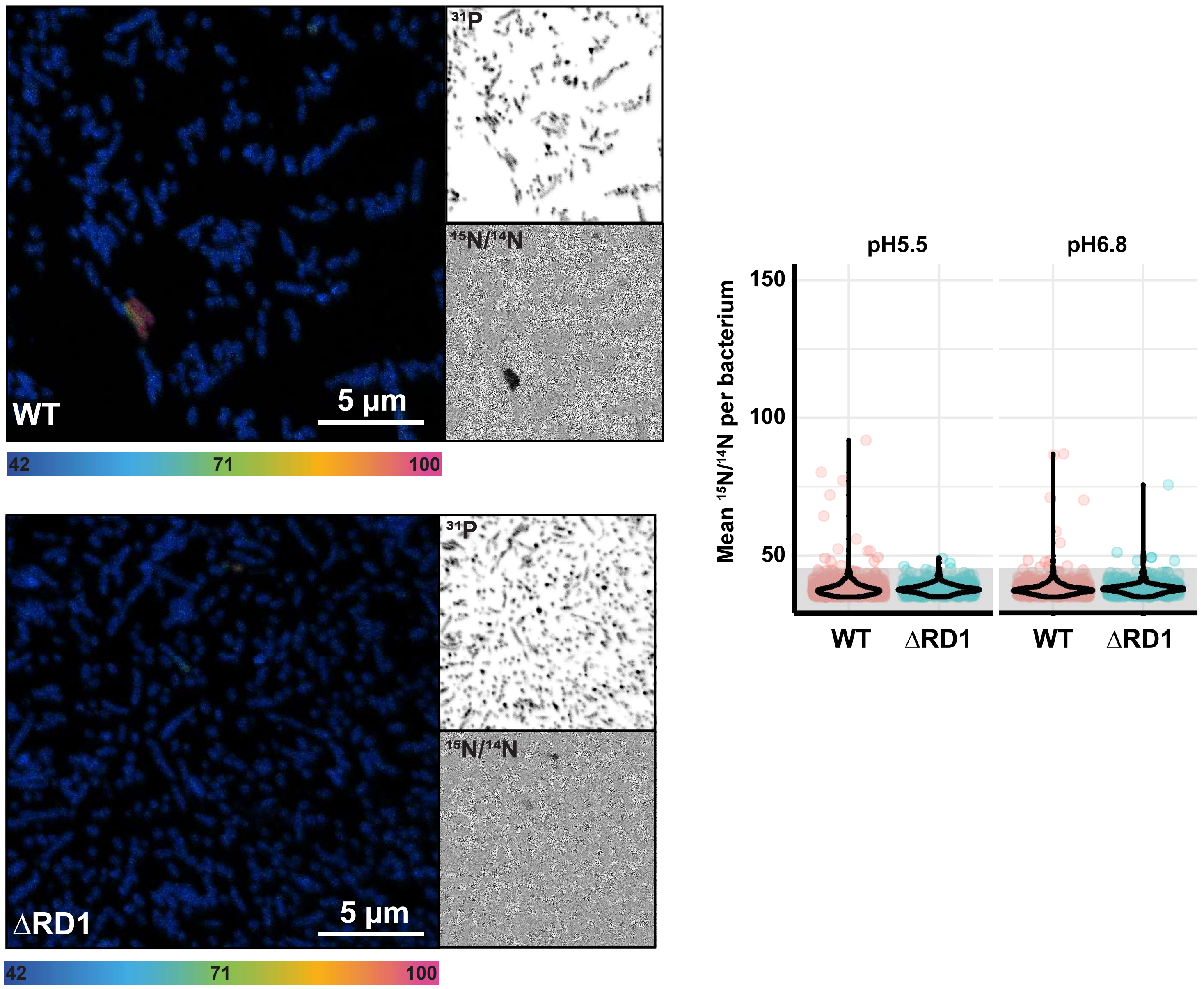
31P
15N/14N
WT
5 µm
42
71
100
ΔRD1
5 µm
42
71
100
pH5.5
pH6.8
Mean 15N/14N per bacterium
150
100
50
WT
ΔRD1
WT
ΔRD1

### Figure S5

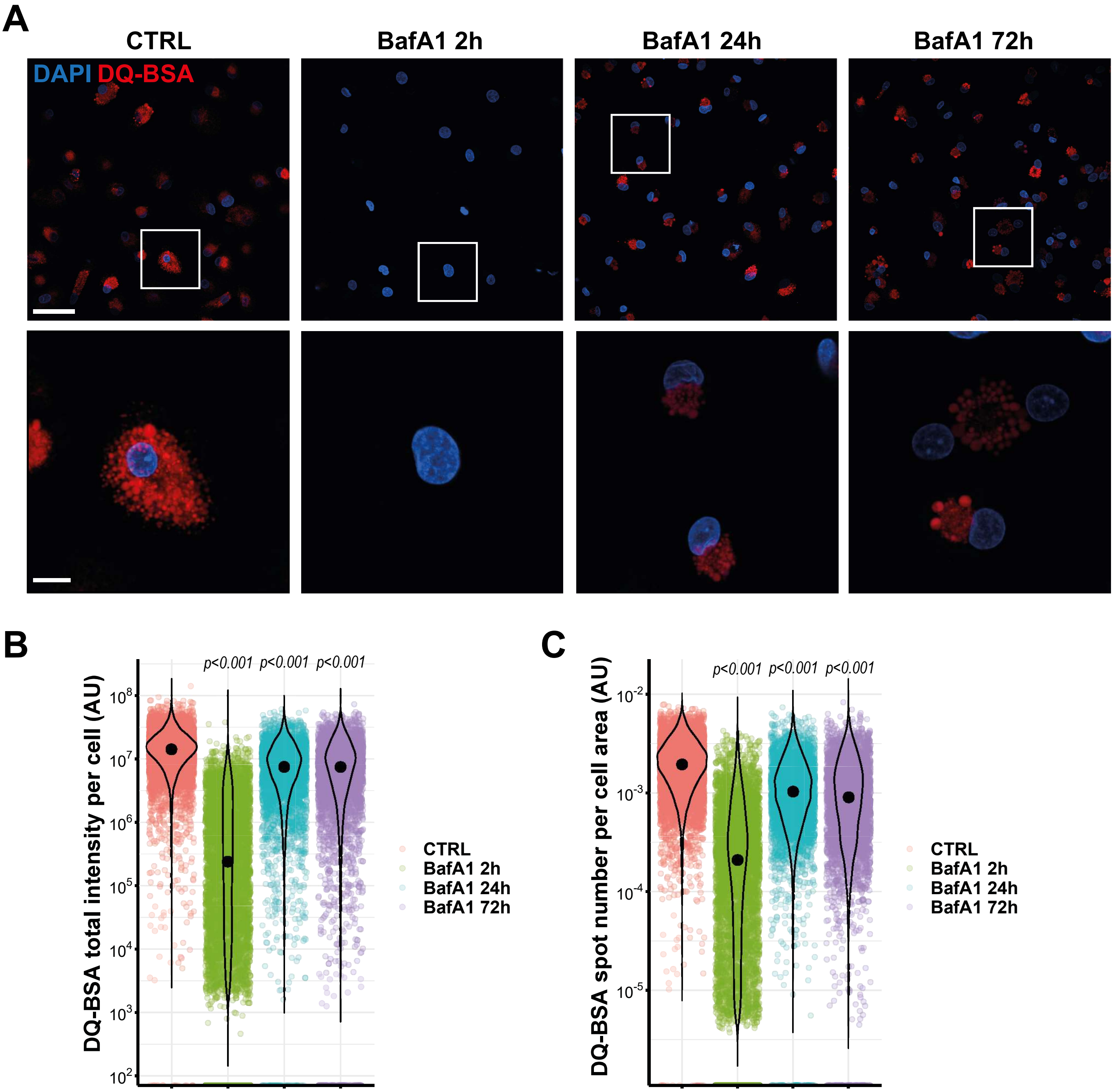
A
CTRL
BafA1 2h
BafA1 24h
BafA1 72h
DAPI DQ-BSA
B
p<0.001 p<0.001 p<0.001
DQ-BSA total intensity per cell (AU)
10^8
10^7
10^6
10^5
10^4
10^3
10^2
CTRL
BafA1 2h
BafA1 24h
BafA1 72h
C
p<0.001 p<0.001 p<0.001
DQ-BSA spot number per cell area (AU)
10^-2
10^-3
10^-4
10^-5
CTRL
BafA1 2h
BafA1 24h
BafA1 72h

### Figure S6

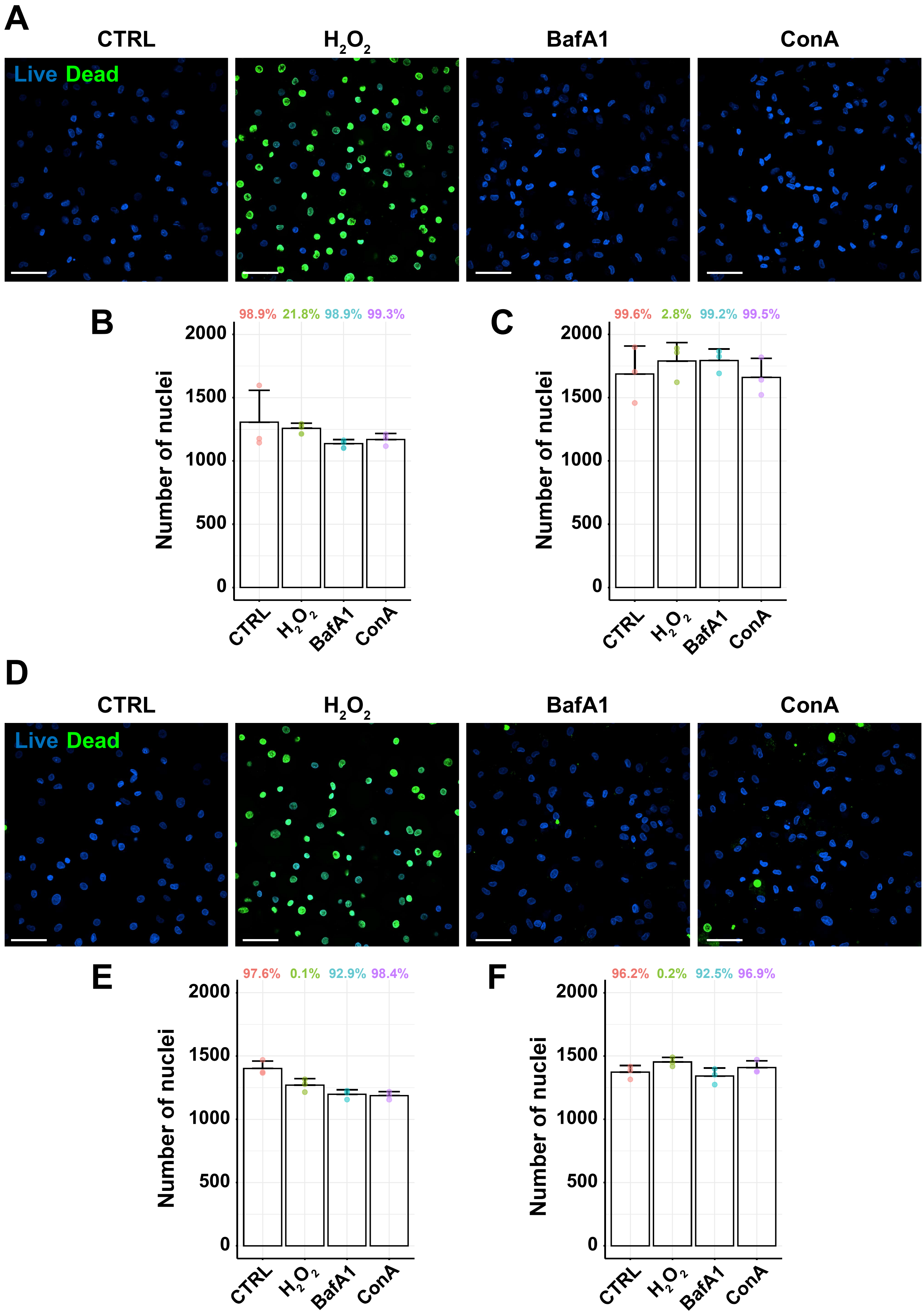
A
CTRL
H2O2
BafA1
ConA
Live Dead
B
98.9% 21.8% 98.9% 99.3%
Number of nuclei
2000
1500
1000
500
0
CTRL
H2O2
BafA1
ConA
C
99.6% 2.8% 99.2% 99.5%
Number of nuclei
2000
1500
1000
500
0
CTRL
H2O2
BafA1
ConA
D
CTRL
H2O2
BafA1
ConA
Live Dead
E
97.6% 0.1% 92.9% 98.4%
Number of nuclei
2000
1500
1000
500
0
CTRL
H2O2
BafA1
ConA
F
96.2% 0.2% 92.5% 96.9%
Number of nuclei
2000
1500
1000
500
0
CTRL
H2O2
BafA1
ConA

### Figure S7

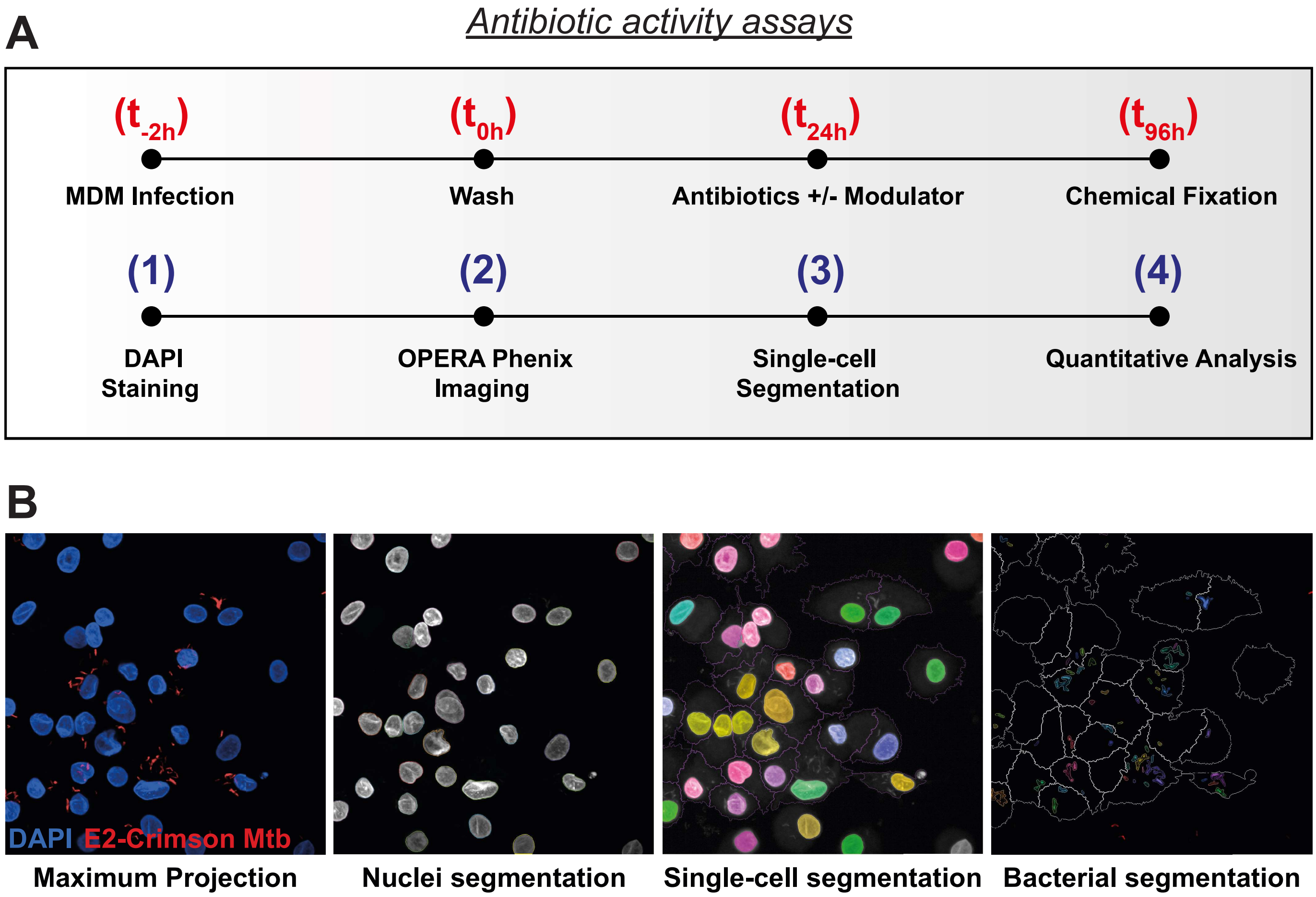
A
Antibiotic activity assays
(t-2h)
MDM Infection
(t0h)
Wash
(t24h)
Antibiotics +/- Modulator
(t96h)
Chemical Fixation
(1)
DAPI Staining
(2)
OPERA Phenix Imaging
(3)
Single-cell Segmentation
(4)
Quantitative Analysis
B
DAPI E2-Crimson Mtb
Maximum Projection
Nuclei segmentation
Single-cell segmentation
Bacterial segmentation

### Figure S8

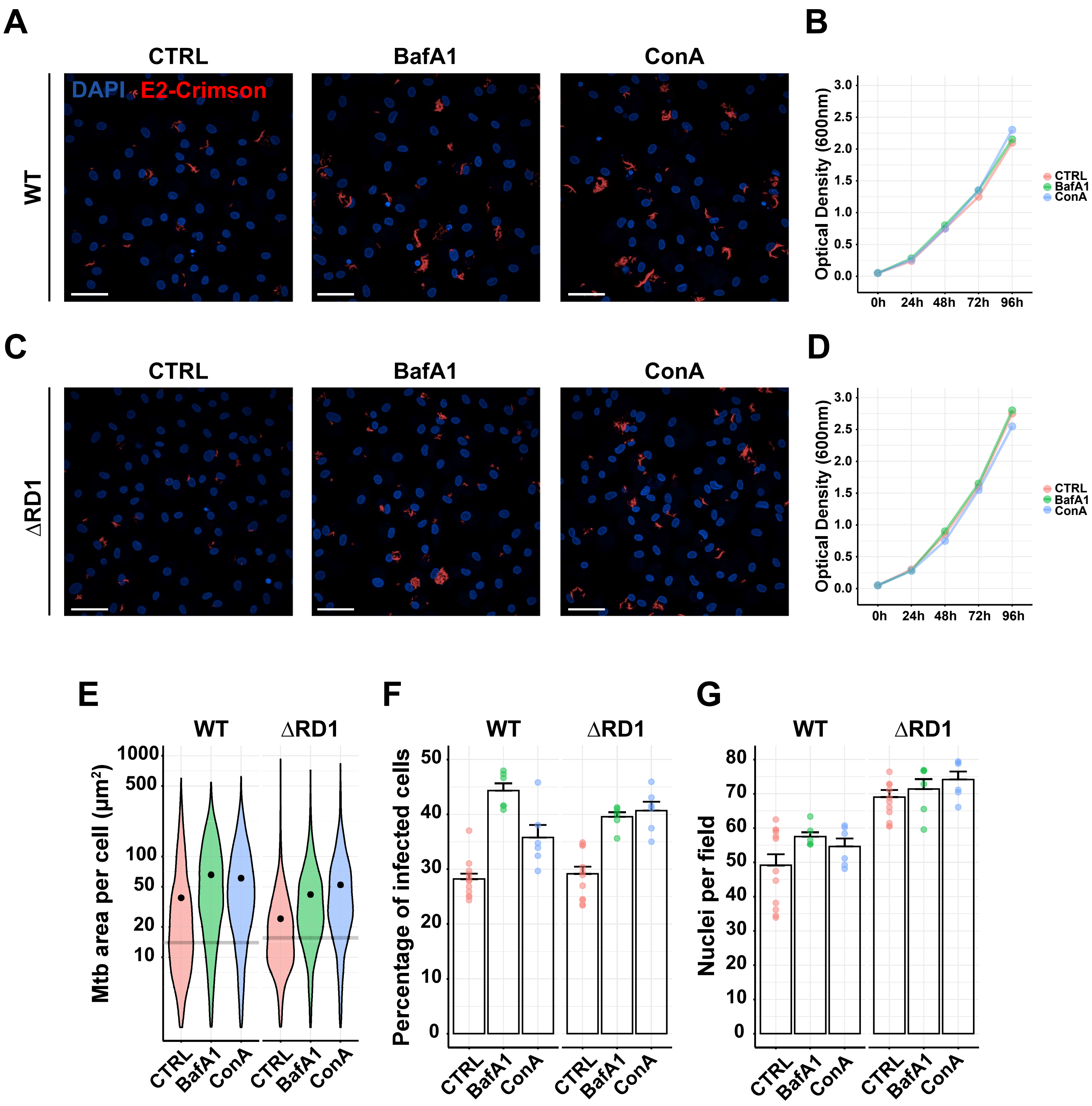
A
CTRL
BafA1
ConA
WT
DAPI E2-Crimson
B
Optical Density (600nm)
0h
24h
48h
72h
96h
CTRL
BafA1
ConA
C
ΔRD1
D
E
Mtb area per cell (µm²)
F
Percentage of infected cells
G
Nuclei per field

### Figure S9

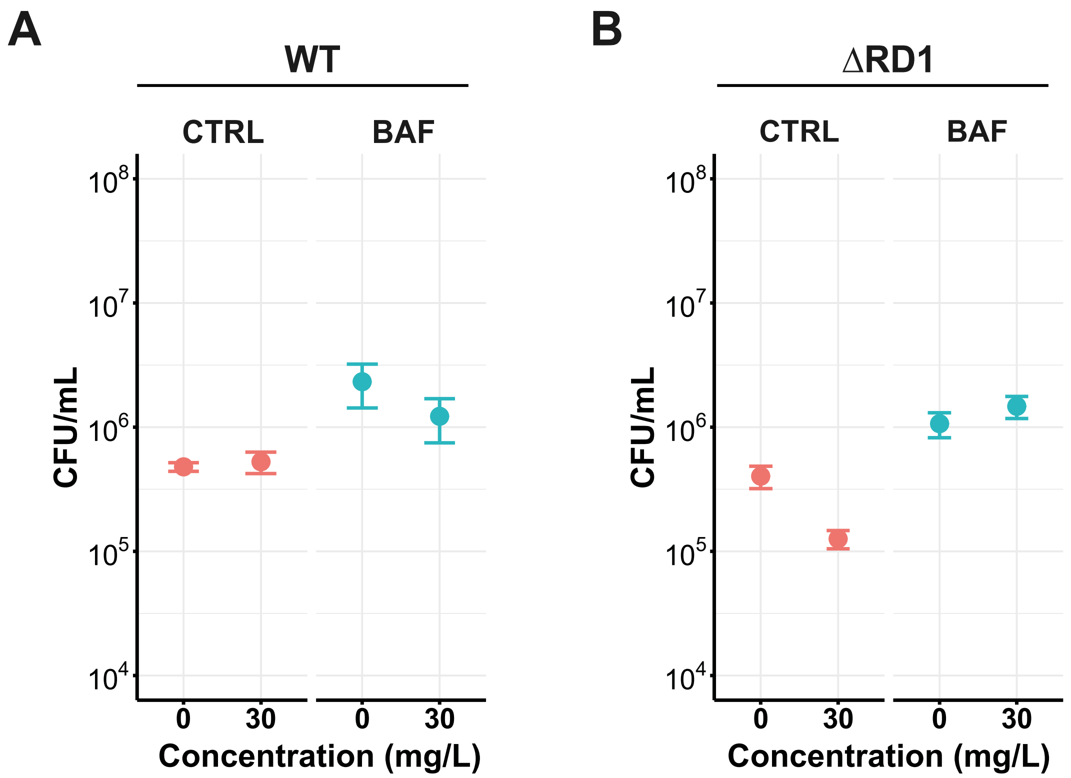

A
WT
CTRL
BAF
CFU/mL
10^8
10^7
10^6
10^5
10^4
0
30
0
30
Concentration (mg/L)
B
ΔRD1
CTRL
BAF
CFU/mL
10^8
10^7
10^6
10^5
10^4
0
30
0
30
Concentration (mg/L)

### Figure S10

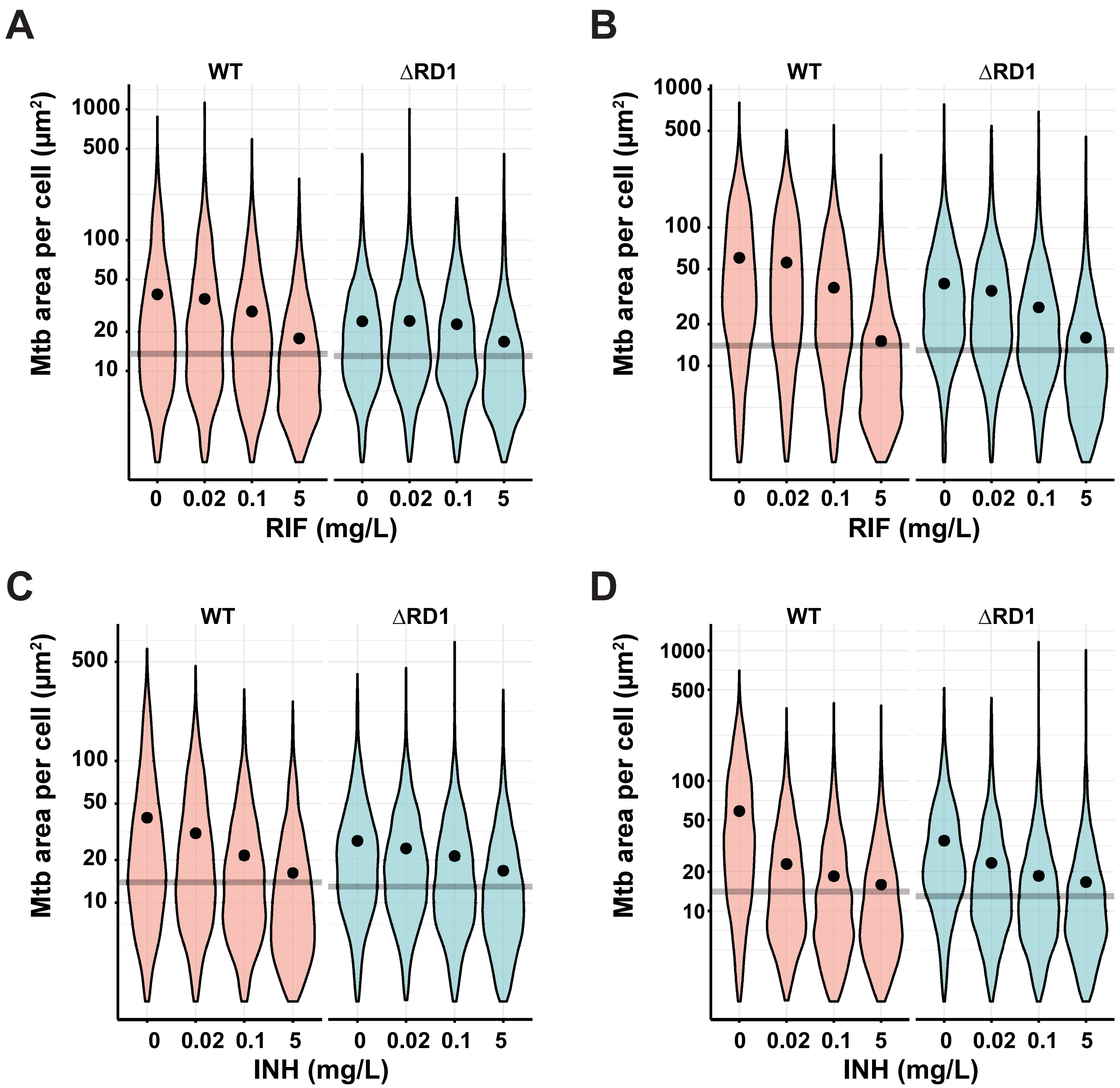

A
WT
ΔRD1
Mtb area per cell (μm²)
1000
500
100
50
20
10
0
0.02
0.1
5
RIF (mg/L)
B
C
INH (mg/L)
D

### Figure S11

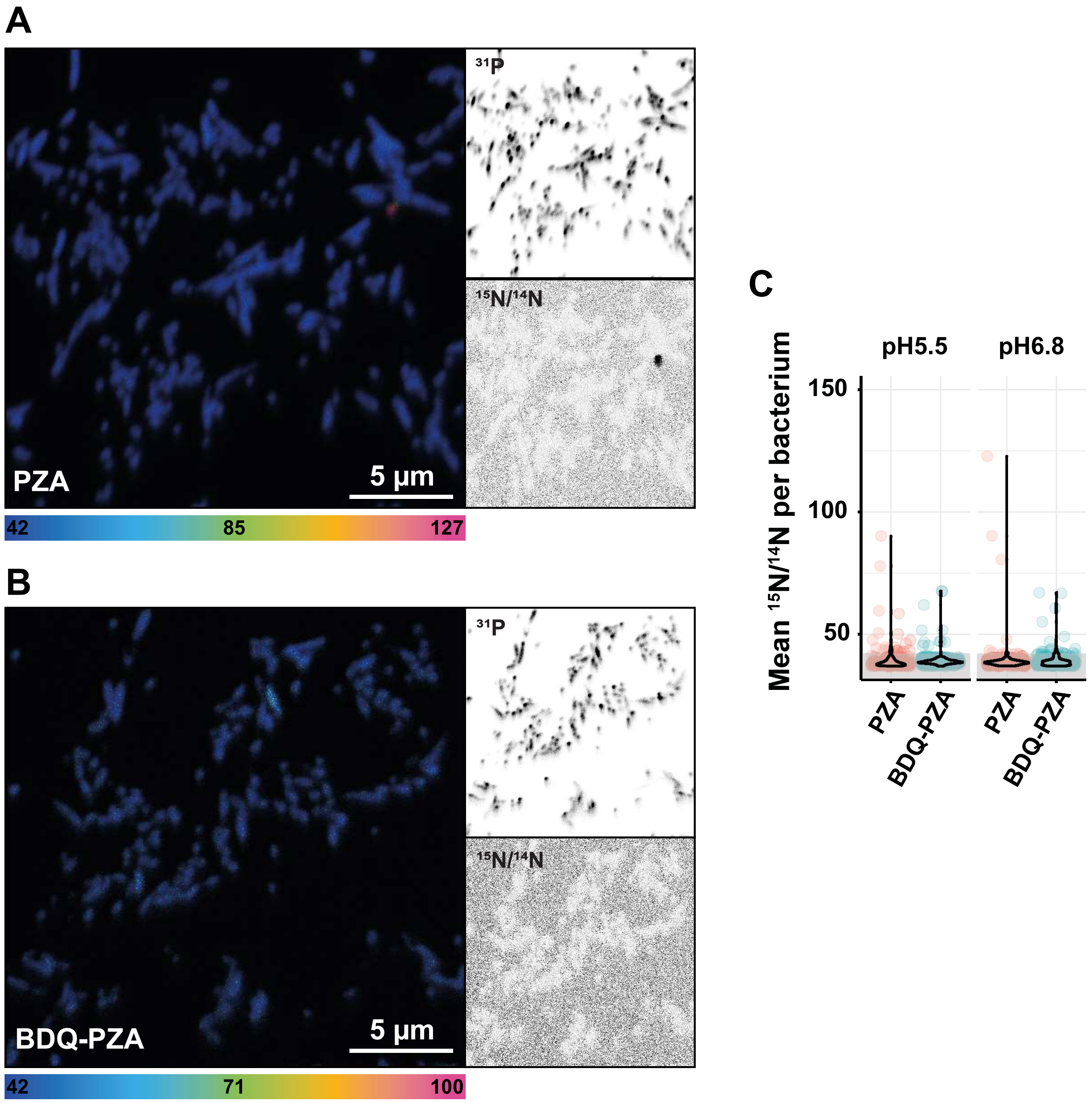
A
B
C
31P
15N/14N
PZA
BDQ-PZA
5 µm
42
85
127
71
100
pH5.5
pH6.8
Mean 15N/14N per bacterium
150
100
50
PZA
BDQ-PZA
